# The Opto-inflammasome in zebrafish: a tool to study cell and tissue responses to speck formation and cell death

**DOI:** 10.1101/2022.10.19.512883

**Authors:** Eva Hasel de Carvalho, Shivani Dharmadhikari, Kateryna Shkarina, Jingwei Rachel Xiong, Bruno Reversade, Petr Broz, Maria Leptin

## Abstract

The inflammasome is a conserved system for the intracellular detection of danger or pathogen signals. By forming a large intracellular multiprotein signaling platform, it activates downstream effectors that initiate a rapid necrotic programmed cell death (PCD) termed pyroptosis and activation and secretion of pro-inflammatory cytokines to warn and activate surrounding cells. However, inflammasome activation is difficult to control on a single-cell level using canonical triggers. We constructed Opto-Asc, a light-responsive form of the inflammasome adaptor protein ASC (Apoptosis-Associated Speck-Like Protein Containing a CARD) which allows tight control of inflammasome formation *in vivo*.

We introduced a cassette of this construct under the control of a heat shock element into zebrafish in which we can now induce Asc inflammasome (speck) formation in single cells of the skin.

We find that cell death resulting from Asc speck formation is morphological distinct from apoptosis in periderm but not in basal cells. Asc induced PCD in can lead to apical or basal extrusion from the periderm. The apical extrusion in periderm cells depends on Caspb but and triggers a strong Ca^2+^ signaling response in nearby cells.

## Introduction

The innate immune system plays a crucial role in early recognition and eradication of potentially dangerous microorganisms. Inflammasomes are among the best characterized components of innate antimicrobial defense in vertebrates. Inflammasome activation is induced following recognition of pathogen-associated molecular patterns (PAMPs) or danger-associated molecular patterns (DAMPs) by intracellular pattern-recognition receptors, such as NOD-like receptors (NLRs) and AIM2-like receptors (ALRs). This leads to the formation of the large multiprotein platforms, which often contain the adaptor molecule ASC (apoptosis-associated speck-like protein containing a CARD), that in turn recruit and activate proinflammatory caspases (Martinon, Burns and Tschopp, 2002). These caspases cleave and thereby activate downstream effector molecules, including the pore-forming Gasdermins (GSDMs) and pro-inflammatory cytokines such as interleukin-1β (IL-1β) and interleukin-18 (IL-18), which are then released through the GSDM pores (Evavold *et al*., 2018; Heilig *et al*., 2018)

Inflammasome pathway components are expressed in immune and non-immune cells, but have been studied extensively mostly in cultured cells or bone marrow derived macrophages (Tweedell, Malireddi and Kanneganti, 2020). Recent studies also highlight their involvement in antimicrobial defense of epithelial tissues (Santana *et al*., 2016; Churchill, Mitchell and Rauch, 2022).

Epithelia are the first point of contact between the host organism and microbial invaders. Although the role of inflammasome components can be studied *in vivo* using epithelial infection models, little is known about the direct effects of inflammasome formation on cells and their neighbors in the context of the live tissue. Thus, analysis of this dynamic and rapid process would benefit from precise spatial control of inflammasome formation and live imaging of the ensuing physiological events.

Zebrafish are well suited to study innate and adaptive immunity *in vivo* (Novoa and Figueras, 2012; Renshaw and Trede, 2012). Their fast development, their transparency in early stages and a multitude of fluorescent reporter lines offer ideal conditions for real-time visualization of cellular processes in tissues. Some of the genes encoding core components of innate immune signaling pathways are highly conserved in the zebrafish (Stein *et al*., 2007), while others, like those encoding the fish-specific NLR proteins, are more divergent (Howe *et al*., 2016). The adaptor protein ASC is among the most highly conserved inflammasome components in all vertebrates. ASCconsists of a pyrin domain (PYD) and a caspase recruitment domain (CARD) connected by a flexible linker. Once activated, ASCsc undergoes prion-like oligomerization, concentrating the entire pool of ASC in the cell in one spot to form a dense fibrous structure termed the ASCsc-speck (Dick et al., 2016).

In zebrafish, Asc is expressed both in epithelial and immune cells, such as macrophages and neutrophils (Kuri et al., 2017). The larval zebrafish skin consists of two cell layers. The outer cell layer, periderm, consists of keratinocytes which show characteristic actin ridges and are tightly connected, the underlying layer consists of basal cells which in adult zebrafish harbor a stem cell pool to replace dying cell in the skin (Lee, Asharani and Carney, 2014). During larval development the periderm is gradually replaced and until then grows by divisions and asymmetric fission (Chan *et al*., 2022).A functional inflammasome can be induced in both layers of the larval zebrafish skin by over-expressing Asc-mKate2, leading to immediate cell death of periderm and basal cells, (Kuri et al., 2017).

Similarly to mammals, zebrafish caspases are recruited to Asc speck (Kuri *et al*., 2017). The zebrafish genome encodes three inflammatory caspases which contain a PYD domain rather than the CARD found in other vertebrates, to recruit the downstream, caspases, Caspa, Caspb (Masumoto, Zhou, Felicia F. Chen, *et al*., 2003) and casp19b (Spead *et al*., 2018) to the Asc speck. The functional homologues of mammalian caspase-1 in zebrafish, Caspa and Caspb, are both able to induce cell death in zebrafish (Kuri *et al*., 2017; Shkarina *et al*., 2022), and GSDMD cleavage and pyroptosis in human cells (Shkarina *et al*., 2022). Caspb has been shown to cleave the zebrafish Gasdermins, Gsdmea and Gsdmeb, in vitro (Chen *et al*., 2021), while caspa has been shown to co-localize with Asc-specks (Masumoto, Zhou, Felicia F Chen, *et al*., 2003; Kuri *et al*., 2017)

Upon cleavage, GSDMs assemble into pores in the plasma membrane and lead to the release of the cytosol including cytokines from the cell (Kayagaki *et al*., 2015; Shi *et al*., 2015; Liu *et al*., 2016; Sborgi *et al*., 2016), a hallmark of pyroptotic cell death (Rühl and Broz, 2021). In the absence of GSDM D, mammalian caspase-1 activates apoptosis as a default pathway (Heilig *et al*., 2020; Tsuchiya *et al*., 2021), and the same was found for ASC-containing inflammasomes in the absence of caspase-1 (Sagulenko *et al*., 2013; Kitazawa *et al*., 2017; Lee *et al*., 2018). Apoptosis is morphologically distinct from pyroptosis and characterized by preservation of membrane integrity, characteristic membrane blebbing and progressive fragmentation of the cell, in contrast to pyroptosis, which most commonly leads to a characteristic cell swelling.

We have previously developed optogenetic variants for human and zebrafish caspases which can be activated by light-induced oligomerization of Cry2olig, a photosensitive protein that undergoes rapid homo-oligomerization in response to blue light (Taslimi *et al*., 2014; Shkarina *et al*., 2022). This enabled us to selectively induce multiple forms of programmed cell death (PCD) in different cell types in vitro, organoids and in the live zebrafish larvae. We observed that zebrafish periderm cells are apically extruded from their surrounding epithelium upon stimulation of the inflammatory caspases but are basally extruded after activation of the apoptotic caspase-8 (Shkarina *et al*., 2022).

To further study inflammasome formation and the resulting cell death in the zebrafish skin upstream of inflammatory caspases, we have now generated an optogenetic variant of zebrafish Asc (Opto-Asc), which efficiently induces speck formation in single cells in both layers of the larval skin. Opto-Asc specks cause inflammatory cell death followed by apical or basal extrusion in periderm cells, which is morphologically distinct from apoptosis. Asc-speck induced cell death is dependent on caspa and caspb but we could not confirm the role of gasdermins in this process, we therefore refrain from calling it pyroptosis. We show that Opto-Asc specks efficiently trigger cell death in the periderm via inflammatory apical or basal extrusion.

## Results

### Optogenetic activation of Asc-oligomerization

The adaptor protein Asc is a core component of the canonical inflammasome and is well conserved in vertebrates. Following interaction with the activated pattern recognition receptors, it forms a large signaling complex (Asc speck) which recruits and activates proinflammatory caspases. In the past, *in vivo* induction of inflammasome assembly has been achieved only via overexpression of Asc or Nlrs (Kuri *et al*., 2017), or by treatment with proinflammatory chemicals like CuSO_4_(Kuri *et al*., 2017) or pathogens (Tyrkalska *et al*., 2016; Li *et al*., 2018; Forn-Cuní, Meijer and Varela, 2019). However, chemical and bacterial treatments affect the entire organism, which complicates targeted and controlled studies of cellular and tissue response. In contrast, the over-expression of Asc efficiently triggers speck formation and cell death, but does not allow targeting these processes in pre-determined cells. To overcome these limitations and to enable specific induction of inflammasome assembly with spatial and temporal control, we tagged zebrafish Asc with Cry2olig to produce “Opto-Asc”. Cry2olig is a protein module that, upon exposure to blue light, reversibly oligomerizes within seconds (Taslimi *et al*., 2014). We fused the mCherry-Cry2olig domain to the N-terminal Pyd domain of Asc and placed this cassette under control of a heat shock responsive element (HSE) (Bajoghli *et al*., 2004), to be able to selectively induce expression only in experimental larvae. We generated stable zebrafish lines for the plasmid shown in Fig. 1A using Tol2-based random genome integration. The plasmid contains a screening cassette in which tagRFP is expressed under the heart specific myosin light chain 2 promoter (cmlc2). Larvae were screened for red hearts at 2.5 days post fertilization (dpf). Opto-Asc larvae were heat-shocked to induce expression and shielded from light to prevent uncontrolled Opto-Asc activation. The photoactivation and imaging experiments were performed at 3dpf as shown in Fig. 1B.

**Figure 1:**
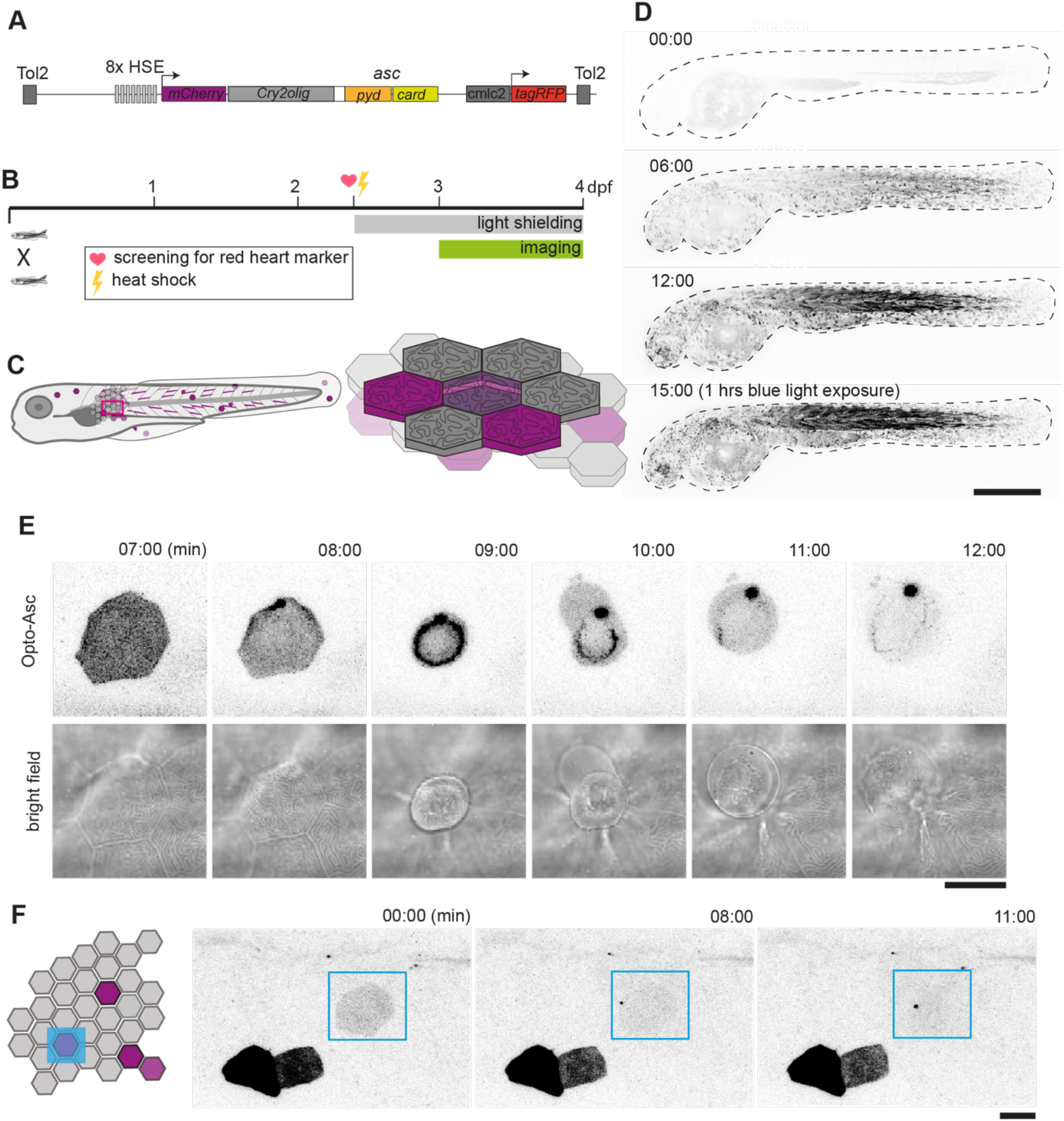
Optogenetic tools for Asc-dependent inflammasome formation. **A**. The optogenetic construct (Opto-Asc) consists of a heat shock element placed upstream of a cassette containing the sequences for *mCherry*, the photoactuator *Cry2olig* domain and *asc*, followed by a cassette containing the “red heart” marker cmlc2:*tagRFP*. The two cassettes are placed between Tol2 sites for insertion into the genome. **B**. Experimental set-up: Progeny from the transgenic lines are first screened for the expression of the red heart marker at 2 dpf. Positive larvae expressing cmlc2:*tagRFP* are heat-shocked and kept in dark conditions to prevent spontaneous opto-Asc activation throughout the experiment. Imaging is performed at 3 dpf. **C**. Schematic depicting the stochastic expression of Opto-Asc. Left: overview of larva; right: diagram of the epidermis with the periderm on top (dark colour) and basal cell layer below (light colour). **D**. Time lapse images of 3dpf larva expressing Opto-Asc. Expression of Opto-Asc becomes detectable at 6 hours post heat shock. The frame rate is 15 min, time points are hours after heat shock, scale bar is 200 µm. **E-F**. Example of Opto-Asc forming specks in the epithelial layer of 3dpf larva expressing Opto-Asc. Scale bars are 20 µm. **E**. Time-lapse imaging after 488nm laser illumination of the periderm cells. Top row: Asc-expressing cell forming a speck (t=8 mins). Bottom row: morphology of the dying cell in bright field. Within a minute of speck formation, the cell changes morphology and is extruded. All bright field images are at the plane of the periderm cells; fluorescent images are z-projections 30 planes (z=1 µm) **F**. Local activation of Opto-Asc in a single cell. Diagram of periderm showing four cells expressing Opto-Asc and region of optogenetic activation (blue square). The cell with the lowest expression of Opto-Asc was illuminated by 2-photon laser. Only this cell forms a speck.

The expression of Opto-Asc became detectable around 3 h after the heat shock and reached a maximum at around 12 h, leading to the mosaic expression of the Opto-Asc in single cells of all tissues Skin cells of different layers can be identified by their distinct morphology: periderm cells, the outer layer of skin cells, have sharp boundaries and a characteristic pattern of actin ridges in the apical cortex. The underlying basal cells are smaller and have less well-defined boundaries (Fig.1C). In absence of blue light stimulation, Opto-Asc did not spontaneously form specks, but did so efficiently when stimulated with the 488 nm laser (Fig. 1D).

The exposure of cells expressing Opto-Asc to 488nm laser light led to the formation of a speck within minutes followed by immediate death of the cell in both epidermal layers (Fig. 1E). In the peridem cells were usually apically extruded from the within minutes after the appearance of the speck. Even when small regions were selectively illuminated with the 488nm confocal laser, the specks also appeared in regions outside the illuminated area, and sometimes even in the entire larva, likely due to the out-of-plane Cry2olig activation and confocal light diffusion, and this could be avoided by using a 2-photon laser, in which case which Opto-Asc oligomerization could be induced in single cells without affecting surrounding cells (Fig.1F).

In muscle cells, illumination induced the formation of multiple stable oligomers of Opto-Asc without inducing cell death (Supplementary Fig 1A), as also previously shown for Asc-mKate2 specks (Kuri *et al*., 2017). These Opto-Asc specks remained stable and did not disperse after light exposure was terminated (for how long?). As control for the contribution of the Asc moiety of the construct to oligomer formation we injected a mutant variant of Cry2olig OptoR489E-Asc, which formed no oligomers at all and did not induce cell death (Supplementary Figure 1B). Additionally, we also tested a C-terminal fusion of Cry2olig to Asc, which also formed light-induced oligomers but did not trigger the formation of large specks or cell death (data not shown). This indicated that the Asc moiety is not functional in this fusion protein, suggesting that the functional induction of speck formation by Opto-Asc in zebrafish requires oligomerization to be triggered close to the Pyd domains which have to interact to form the initial filament (Dick *et al*., 2016) or that zebrafish ASC requires some post-translational modification that is hampered by the Cry2olig domain.

Kuri et al. showed that Asc specks could also be induced by overexpression of the Pyd-like domain of one of the zebrafish NLR family members (si:dkeyp-4c4.1), most similar to mammalian NLRP1. To induce speck formation upstream of Asc we generated optogenetic variants of the full-length version as well as the Pyd domain of the NLR. However, the expression of mCherry-Cy2olig-Nlr-FL, mCherry-Cry2olig-Nlr-Pyd or Nlr-Pyd-Cry2olig-mCherry did not efficiently trigger speck formation in a time frame that was comparable to Opto-Asc (data not shown).

### Optimization of heat shock and light exposure conditions for efficient Opto-Asc speck induction

During our initial experiments, we observed variability in the efficiency of speck formation both between different larvae and between cells within a single larva. Thus, to identify conditions for efficient induction of speck formation, we titrated both the heat shock duration to maximize the expression of the construct and the laser illumination parameters to optimize the light-induced oligomerization of Cry2olig.

To first determine the optimal heat shock parameters, we crossed the Opto-Asc line to a Krt4:AKT-PH-GFP line, which enabled us to visualize the membrane outlines of skin cells in order to count individual cells (Fig.2A). We then incubated the larvae at 39°C for different periods of time and calculated the fraction of periderm cells expressing detectable levels of Opto-Asc and the overall mean fluorescence intensity (MFI) of a defined region in each larva (Fig.2C). While we found no significant correlation between the number of expressing cells and overall MFI in any of the heat shock schemes (Fig.3D), we detected high expression (>12.000 MFI) and a high percentage of expressing cells (>30 %) in those larvae that had been heat-exposed for 35 min or longer. After longer heat shocks (40 min) the larvae started to show deformations in their somites. We therefore performed all further experiments with 35 min heat shocks and, unless stated otherwise, used larvae expressing Opto-Asc in >30 % of their cells for evaluation.

**Figure 2:**
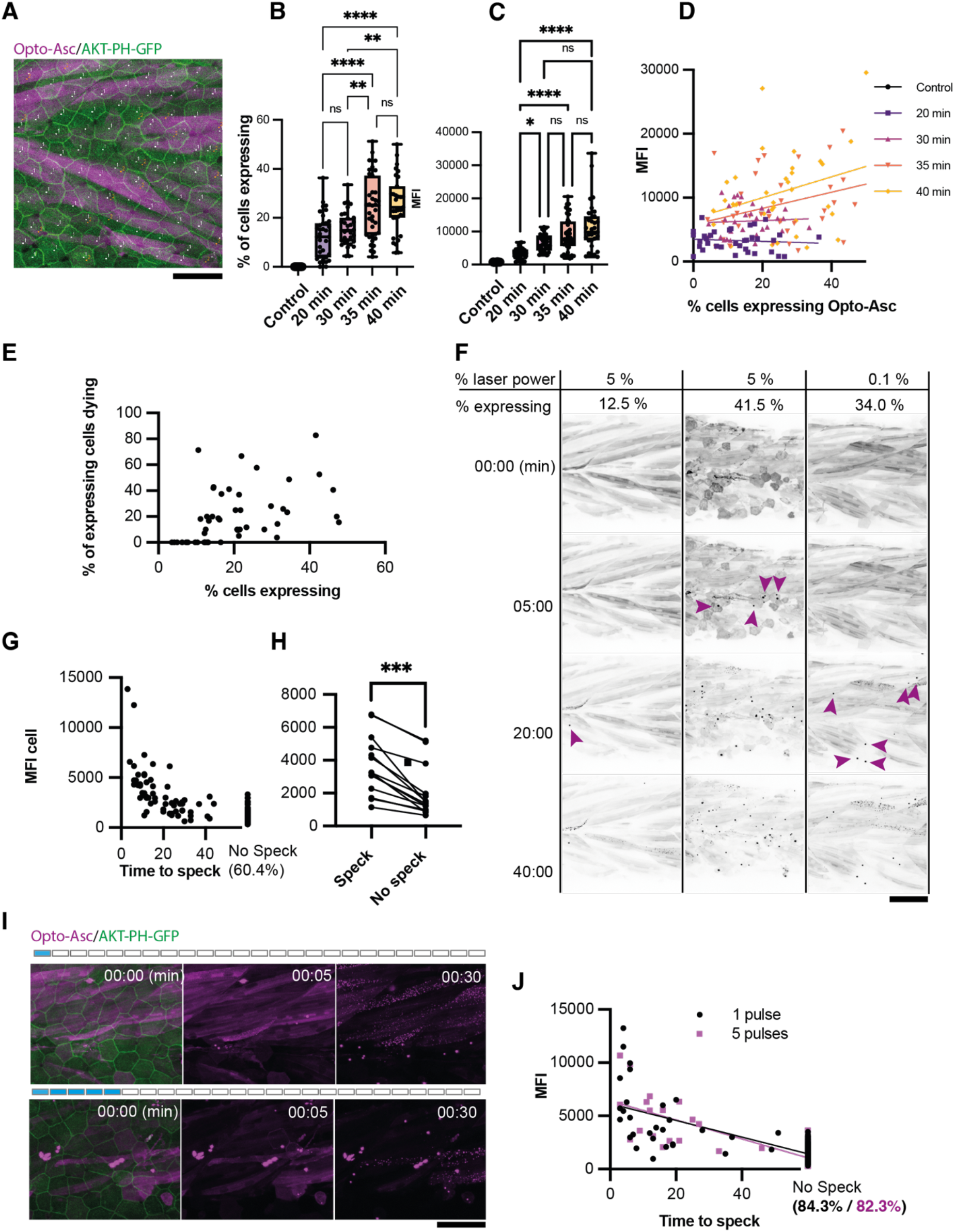
Effect of heat shock and light exposure. **A**. Periderm in a larva expressing Opto-Asc (magenta) and AKT-PH-GFP to mark membranes (green) at 16 hr post heat shock (hrpHS). Cells are individually numbered for subsequent quantification of Asc expression. Opto-Asc in this figure is mainly visible in muscles underlying the epidermis, which are seen in this maximum Z-projection. For quantification of Asc-expressing periderm cells, individual z-planes within the periderm are evaluated. Scale bar: 50 µm **B-D**. Dependence of Opto-Asc expression on heat-shock protocol **B**. Percentage of Opto-Asc expressing cells and **C**. overall mean fluorescence intensity (MFI, measured on max projections) in larvae heat-shocked for different time periods (N>29 for each condition; each dot represents one larva). **D**. Correlation between percentage of Opto-Asc-expressing cells and overall MFI for all larvae for each condition measured in B (N=136/>29 per condition). **E**. Correlation between the percentage of Opto-Asc expressing cells and the number of expressing cells dying within 60 min (N=47 movies; each dot one larva); all movies at 1 min frame rate, 5 % relative laser power. **F**. Examples of larvae expressing Opto-Asc after exposure to 488 laser light at different laser powers. The number of Opto-Asc expressing periderm cells (shown as % positive cells) does not correlate strictly with the level of laser power, nor does the expression level in the underlying muscles. If a sufficient number of periderm cells are positive, specks are formed even at low laser power. First appearance of specks is indicated by magenta arrows. **G**. Correlation between MFI of Opto-Asc in single cells and the time from beginning of 488 laser light exposure to speck formation. Cells were followed for 60 min; 60.4% of the cells had not formed a speck at this time. N=156 cells from 14 different larvae. **H**. Comparison of average MFI between Opto-Asc expressing cells in individual larvae that form and do not form a speck within 60 min (N=15 individual larvae). The values representing the average MFI for speck-forming and non-speck-forming cells for each larva are connected by a line; p<0.001. **I**. Time lapse of Opto-Asc expressing larva exposed to a single or to 5 laser light pulses (1 or 5 frames acquired with 488 laser light). Scale bar is 50 µm. **J**. Correlation between MFI of single cells and onset of speck formation in cells exposed to either 1 (red circles) or 5 (black outlined squares) pulses of 488 laser light. Cells were followed for 60 min. N=164 cells in 9 larvae for 1 pulse, N=86 cells in 5 larvae for 5 pulses.

**Figure 3:**
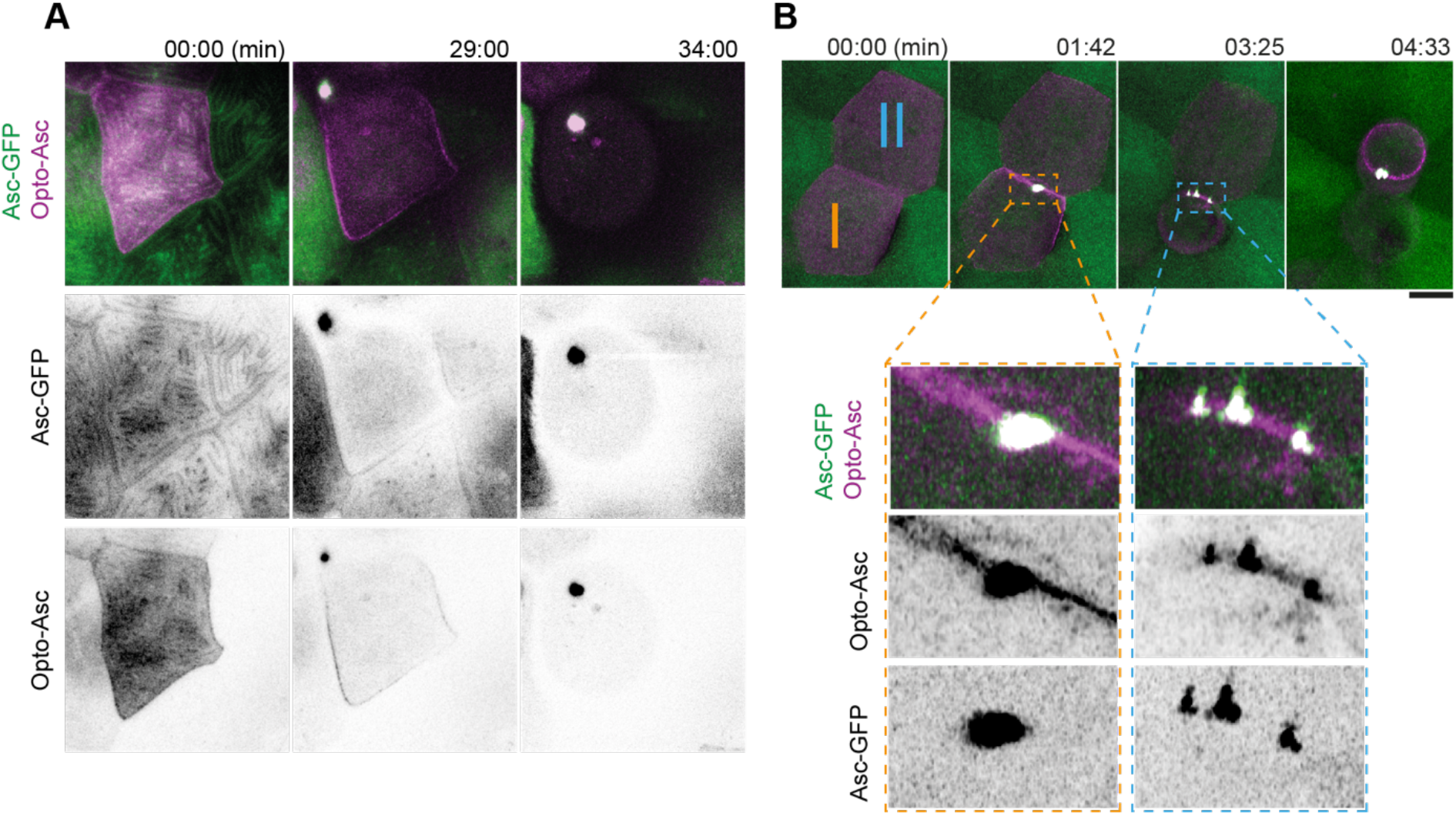
Speck formation dynamics of Opto-Asc and recruitment of endogenous Asc. **A**. Time lapse of speck formation induced by Opto-Asc (magenta) and recruitment of endogenous Asc (green, ubiquitous) in a periderm cells. Scale is 10 µm. **B**. Two neighboring cells (I and II) forming either a single speck (cell I) or multiple specks (cell II) along the cell membrane. The specks coalesce as the cell shrinks. Scale is 2 µm.

To next determine the effect of Opto-Asc expression levels on the efficiency of speck formation, we exposed the heat-shocked larvae to 5% (1.24mWatt/µm2) 488 nm laser light (an intensity normally used to image our GFP reporter lines). After 60 minutes, we counted the percentage of Opto-Asc-positive epithelial cells in the imaged area and the percentage of speck-forming cells among those Opto-Asc positive cells (Fig.2E). We found no correlation between the number of positive cells and the likelihood of speck formation. However, specks only ever formed in larvae in which more than at least 10 % of the cells were Opto-Asc positive, indicating that there is a threshold for Opto-Asc expression below which cells do not form a speck. Figure 2F shows three examples of larvae with different fractions of Opto-Asc expressing cells which had been exposed to 5 % or 1 % 488 nm laser light. Larvae in which few cells expressed Opto-Asc (here 12.5 % of cells) rarely formed specks, even after long exposure times (1 stack/min for 60 min). Larvae with many expressing cells formed specks within less than five minutes, regardless of laser power. In larvae with high Opto-Asc levels, laser power as low as 0.1 % (0.025 mWatt/µm^2^) was sufficient to induce speck formation after several minutes (Fig. 2F).

We also observed an Opto-Asc expression level threshold for speck formation when assessing individual cells. We measured the MFI of single cells in 14 larvae and scored for each cell the time it took to form a speck under repeated blue light exposure for 60 minutes (Fig.2G). While we found no absolute threshold for speck formation, 39.6 % of the cells with Asc levels of >5000 MFI (relative units) formed specks within the period of observation, usually within the first 30 min of exposure. Cells that did not form a speck within 60 min (60.4 %) all had an MFI of < 5000 (relative units). When comparing cells only within individual larvae, we similarly found that cells which did not form specks also had significantly lower opto-Asc expression levels (Fig.2H). We also compared the effects of constant illumination and a pulsed laser exposure, where a pulse is defined as the period of imaging of a single stack (40 µm in 41 steps). We compared the effect of stimulation with one and five pulses (Fig.2I) on the cells with the approximately similar opto-ASC expression levels, which revealed that compared to constant light exposure (e.g. Fig.2H), fewer cells formed a speck after pulsed illumination: 15,7% for five pulses (average MFI: 1999) and 17,7 % after one pulse (average MFI: 2063). The specking response to laser pulses was delayed, sometimes up to 50 min (Fig.2J), similar to the dynamics we see at constant laser exposure (cell death after up to 40 min post first illumination. While for most oligomerization reactions, a single initial laser pulse was sufficient to trigger speck formation, a longer light exposure increased the fraction of cells in which Cry2olig oligomerization was triggered (Fig.2G).

Based on all of these observations, we chose the following conditions for our further experiments: after 35 min HS we select larvae based on the percentage of expressing cells (above 20%) and exposed them to 5% relative 488 nm laser intensity.

### Opto-Asc speck appearance and recruitment of endogenous Asc

We have previously shown that specks induced by Asc-mKate2 efficiently recruited endogenous Asc in the asc:asc-GFP line, where the open reading frame of endogenous Asc was tagged with GFP (Kuri *et al*., 2017). To test if Opto-Asc behaves in the same way as Asc-mKate2, we assessed endogenous ASC and opto-ASC co-localization in single cells in larvae expressing endogenous Asc (ubiquitous expression) and Opto-Asc (mosaic expression) using Airyscan high resolution microscopy (Fig.3A). Endogenous Asc and Opto-Asc colocalized in the cortical actin ridges, and during a speck formation they co-aggregated with the same dynamics (Supplementary Fig. 2A and B). Like Asc-mKate, Opto-Asc was present throughout the entire speck together with endogenous Asc (Supplementary Fig.2 C and D).

Opto-Asc specks were on average smaller than Asc-mKate2 specks (Supplementary Fig. 2E), and unlike Asc-mKate2, Opto-Asc occasionally formed multiple specks which often subsequently fused as the cell started to die. As an example of this, Fig.3B shows two neighboring cells with similar Opto-Asc expression forming specks after light exposure. Both cells recruited Asc to their membrane but while cell I formed a single speck, cell II formed multiple irregularly shaped specks.

### Role of endogenous Asc and its domains in Opto-Asc-induced speck formation and cell death

All experiments reported so far were done in fish that had normal, endogenous Asc in addition to the Opto-Asc. Since Opto-Asc recruites endogenous Asc, we do not know whether cell death is triggered by endogenous or Opto-Asc. To test this, we used an Asc loss-of-function mutant, Asc_Δ2/Δ2_, in which the endogenous ASC has a CRISPR/Cas9 induced frameshift mutation that introduces a premature stop codon p.Phe13*to create a protein-null allele. To determine the ability of two different ASC domains, PYD and CARD, to induce speck formation and cell death we egneratd Opto-PYD and Opto-CARD (both N-terminally fused to mCherry-Cry2olig). We injected full-length Opto-Asc, Opto-Pyd and Opto-Card into the Asc_Δ2/Δ2_mutant line. We first tested the ability of Asc-mKate2 to function in this mutant and found that it formed specks and induced cell death at the same level as in the wild-type larvae, showing that Asc-mKate2 is fully functional and able to induce cell death (Fig. 4A).

**Figure 4:**
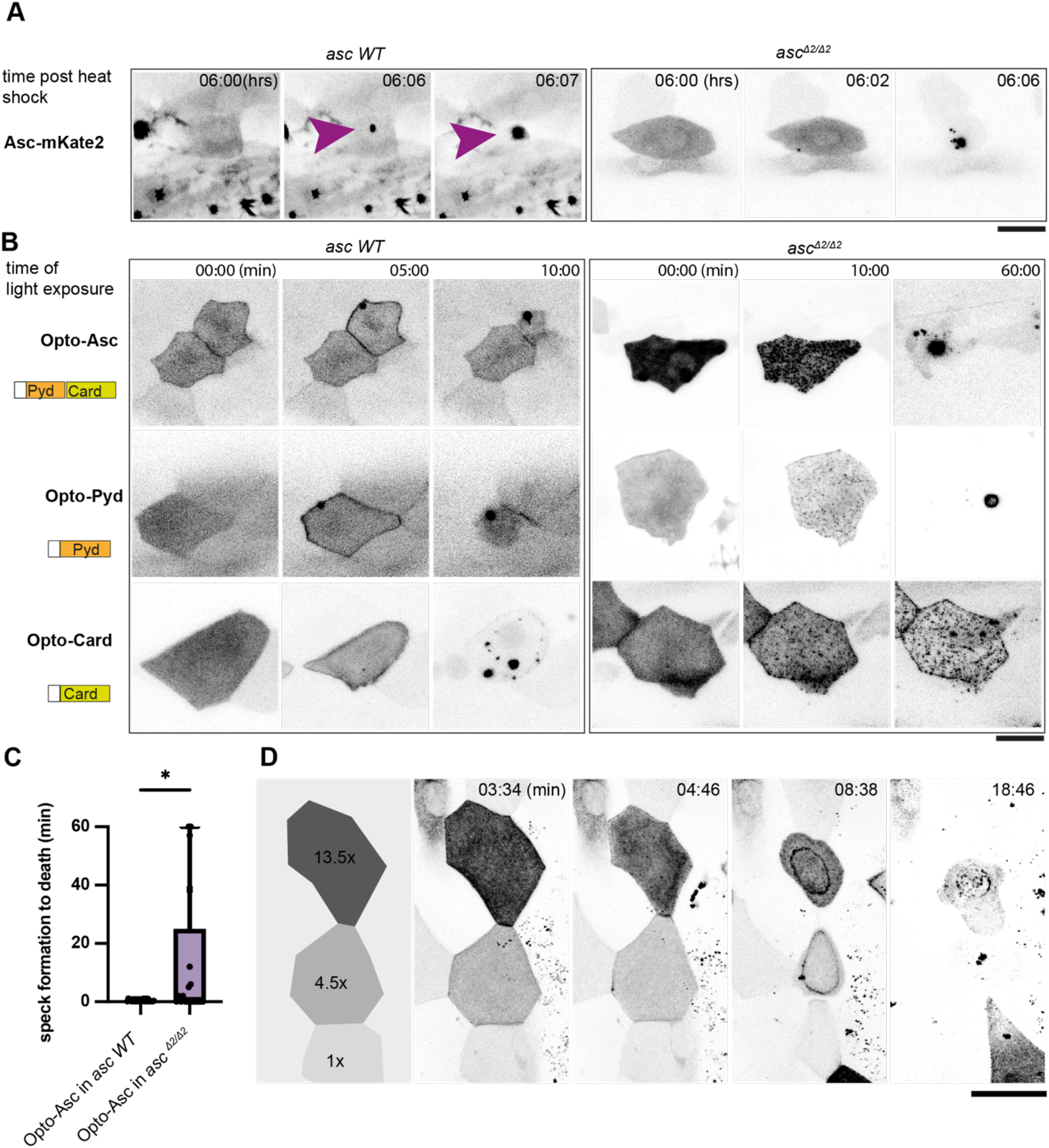
Opto-Asc speck formation and dependence on endogenous ASC. **A, B**. Speck formation induced by Asc-mKate2, Opto-Asc, Opto-Pyd and Opto-Card in wildtype larvae and *asc*_*Δ2/Δ2*_ larvae. Red frames indicate frames in which cells are dying. Scale bar is 20 µm. **A**. Time for Asc-induced specks is counted from end of heat shock. Purple arrow heads mark the speck in the cell of interest. Specks formed by surrounding cells can be seen as balck spots **B**. Time is in min counted from the last timepoint before visible speck formation. **C**. Time is measured between first detectable appearance of a speck and first morphological sign of cell death (deformation of cell) in wildtype and *asc*_*Δ2/Δ2*_ larvae. *=p<0.05. **D**. Dynamics of speck formation in three cells with different expression levels of Opto-Pyd. Relative intensity levels shown in the diagram on the left are measured in relation to the cell with the lowest level (1x) after background subtraction. Time is measured in minutes; scale bar is 20 µm.

In the wild-type Asc-GFP line, Opto-Asc, Opto-Pyd and Opto-Card induced speck formation within five minutes, whereas showed a delayed speck formation (>2.5 min after first aggregates were observed (N=3) The shrinking of the cell, which we consider an initial sign of cell death upon opto-ASC activation, was seen as soon as the first oligomerization was detectable for all of the constructs.

By contrast, in the Asc_Δ2/Δ2_ knockout line, Opto-Asc, Opto-Pyd and Opto-Card formed multiple smaller aggregates rather than a single large speck, indicating that endogenous Asc may facilitate multiple Opto-Asc to coalesce to form a single speck structure. In this mutant, Opto-Card did not induce cell death, consistent with the finding that inflammatory caspases are recruited via the Pyd domain of Asc (Kuri *et al*., 2017), and therefore, without endogenous Asc, caspases could not be recruited to the Opto-Card construct. By contrast, both Opto-Asc and Opto-PYD induced cell death, but often with a delay when compared to Asc WT cells. (Fig. 4B).

When we measured the time between the first detectable oligomerization and first visible signs of cell death, we found that in some Asc_Δ2/Δ2_ knockout periderm cell death was significantly delayed while in other AscΔ2/Δ2 and WT periderm the death was immediate after the first oligomers were detected (Fig.4C).

These observations imply that the assembly of Asc into a clearly defined speck was not required for downstream effectors to induce cell death. Figure 4D shows the example of three cells expressing different levels of Opto-Pyd, which are normalized relative to the cell with the weakest expression (1x). The cell with the highest level of Opto-Pyd (13.5x) entered cell death before oligomerization was detectable, whereas in cells with intermediate (4.5x) or low (1x) levels Opto-Pyd recruitment to the membrane and oligomerization significantly preceded the appearance of first signs of cell death. This shows that the efficiency of cell death induction is dependent on the amount of Opto-Asc or Opto-Pyd, and that in case of high concentrations of Opto-Asc or Opto-Pyd, the formation of a visible speck is not required for initiation of cell death.

### Cell extrusion after Opto-Asc speck formation

Periderm cells typically respond to inflammasome formation by rapid cell death and apical extrusion (Kuri *et al*., 2017). Thus, we took advantage of opto-Asc-mediated inflammasome formation to characterize the cellular events immediately following speck formation. Figure 5A shows an example of a cell that was extruded from the periderm after speck formation. The cell began to shrink within 30 sec of speck formation while the surrounding cells moved in to fill the space. The dying cell lost its shape and started to form membrane blebs, eventually swelled and was finally extruded from the cell layer. Once completely extruded, the cell appeared round and lysed within minutes, as determined by the uptake of the membrane-impermeable DNA-binding dye DRAQ7 which can only stain the cells following membrane permeabilization (Fig. 5B). Most cells reached the lytic stage in 5-10 min post illumination, although some remained fully extruded without lysing for up to 20 min post speck formation (Fig. 5C).

**Figure 5:**
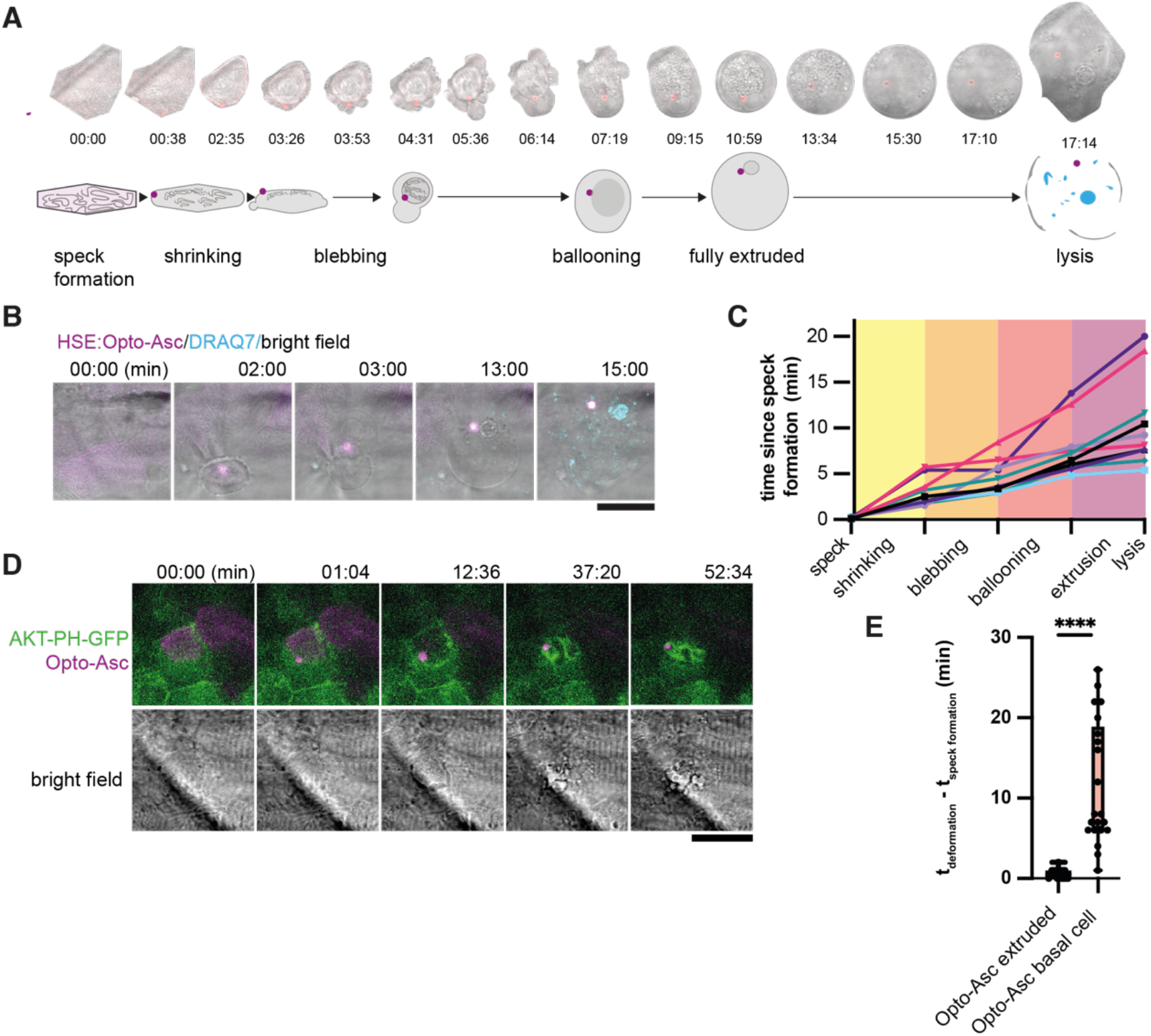
Stages of Asc-speck-induced e extrusion from the periderm. **A**. Example of a single cell from a bright-field movie (top) and a schematic showing stages of periderm cell extrusion and lysis after speck formation (bottom). **B**. DRAQ7 staining of a cell that lyses after the full extrusion from the periderm. Scale bar is 20 µm. **C**. Time lines for 10 periderm cells undergoing the steps of cell extrusion described in A. Time is in minutes. **D**. Basal cell dying after formation of an Opto-Asc-speck (magenta). Cell membrane is labeled by AKT-PH-GFP (green) **E**. Time between Opto-Asc speck formation and the appearance of first morphological signs of cell death (deformation of cell) in periderm and basal cells. ****=p<0.0001

Basal cells in the cell layer underlying the periderm also died following Opto-Asc speck formation, but with a different morphology. They showed extensive fragmentations seen by occurrence of membrane vesicles labeled by Akt-PH-GFP (Fig. 5D). The basal cells also responded more slowly to speck formation than periderm cells and started to change morphology only after ∼5 minutes (Fig. 5E).

### Apical and basal cell extrusion in response to Opto-Asc specks

The fate of the dying cells following Opto-Asc speck formation varied between cells within and between different larvae. Some periderm cells were not extruded apically, as described above, but instead left the periderm on the basal side. Another subset of cells fell in neither of these groups; instead, part of their cell body was extruded apically, rounding up and lysing, whereas another part was extruded basally, morphologically resembling the basally extruded cells. As an example of this, Figure 6A shows cells located near each other in the same larva, responding by apical extrusion (A), basal extrusion (B) or mixed extrusion (M). To see if this variation in phenotype was determined by genetic differences in our fish lines of mixed genetic background (Golden and AB2B2), we injected the Opto-Asc construct into different commonly used laboratory strains (Fig. 6B). In most strains (AB, AB2B2 and Golden) basal extrusion of periderm cells only occurred in a small number (<5 %) of cells per larva, and not in all larvae. However, the WIK (Wild Indian karyotype) strain showed greater variation, as did our experimental line (*Tg(Opto-asc)* x *Tg(Krt4:AKT-PH))* of mixed genetic background (Opto-Asc: Golden and AB2B2, Krt4:AKT-PH: unknown). In these two lines, considerably higher percentages of dying cells were basally extruded (38 % in WIK UPenn and 64 % in *Tg(Opto-asc)*x *Tg(Krt4:AKT-PH)*).

**Figure 6:**
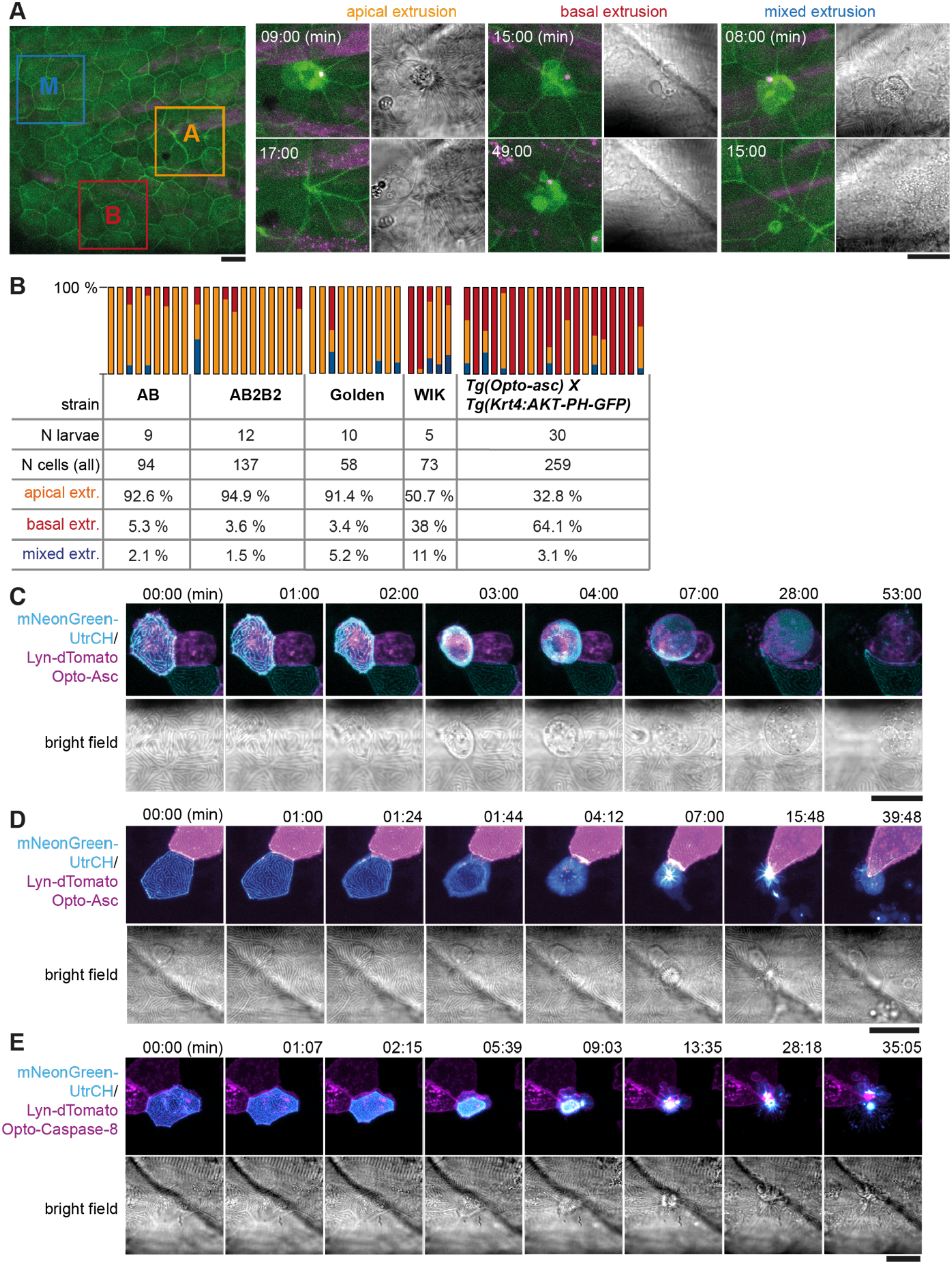
Apical and basal extrusion during Opto-Asc induced cell death. **A**. Examples of responses to Opto-Asc (magenta) speck formation in three periderm cells within the same larva (marked by blue, orange and red squares in the overview image); cell borders marked by AKT-PH-GFP, green. Dying periderm cells can be extruded apically (orange), basally (blue) or in both directions (red). **B**. Percentage of periderm cells extruded either apically, basally or in both directions (mixed) after Opto-Asc speck formation in different *Danio rerio* laboratory strains and transgenic *Tg(Opto-asc X Krt4:Akt-PH-GFP)* larvae. Each bar represents one larva, the y-axis shows what fraction of cells undergoes which type of extrusion. **C-D**. Response of the actin cytoskeleton in cells that are apically (C) or basally (D) extruded after formation of an ASC speck (magenta). Actin is labeled using mosaic expression of mNeonGreen-UtrCH (cyan). The apical cortical actin ridges can also be seen in bright field images. Membranes of cells are mosaically labeled by expression of lyn-tagRFP (also magenta). **E**. Actin response to Opto-Caspase-8-induced apoptosis in periderm cells; actin in the dying cell is in cyan (mNeonGreen-UtrCH). Membranes of cells are mosaically labeled by lyn-tagRFP (also magenta). Scale bars in all images are 20 µm

To characterize the morphological differences between apical and basal cell extrusion, we analyzed the dynamics of the actin cytoskeleton in the dying cells. We visualized actin with an mNeonGreen (mNG)-tagged variant of the calponin homology domain of utrophin (UtrCH) expressed under control of the UAS-promoter (*Tg(UAS:mNG-UtrCH*), which we combined with a Krt4:Gal4 driver and UAS:lyn-tagRFP to label plasma membranes. Larvae carrying these constructs showed stochastic expression of mNeonGreen-UtrCH in periderm cells, which allowed us to follow actin dynamics in single cells. The overall response of the actin cytoskeleton was similar in apically (Fig.6C) and basally (Fig. 6D) extruded cells. As the cell started to shrink after speck formation, the actin ridges rapidly disappeared. Actin then formed a diffuse ring at the periphery of the cell just prior to extrusion. The actin ring contracted on the side of the cell that remained in contact with the epithelium. Basally extruded cells were fragmented once fully internalized and were taken up by surrounding cells or macrophages (Supplementary Fig. 3).

We previously showed that periderm cells dying by opto-zfCaspase-8-induced apoptosis were also basally extruded from the periderm(Shkarina *et al*., 2022). Thus, we analysed the behaviour of their actin cytoskeleton in larvae derived from a cross of *Tg(Opto-caspase-8)* with *Tg(Krt4:Gal4_ UAS:mNG-UtrCH-UAS:Lyn-tagRFP)* and found differences in the apical actin ridges behavior. In contrast to the Opto-Asc-activated cells, apoptotic periderm cells retained the original pattern of actin ridges while the cell was shrinking, with a proportional scaling of the pattern until the cell was fully internalized (approximately 10 minutes) (Fig.6E). While analyzing the behavior of the actin cytoskeleton in the cells surrounding dying cells either after formation of an Asc speck or induction of apoptosis by Opto-Caspase-8, we found that actin was always recruited to the membrane region that was in contact with the dying cell, independent of the cell death phenotype (Supplementary Fig.4). We also analyzed the actin cytoskeleton in basal cells dying in response to Asc-speck formation, which revealed that while the cell was shrinking, its actin was re-localized to the membrane along the borders to neighboring cells (Supplementary Fig. 5A) at the same time as those cells started to migrate to close the gap created by the dying cell (Supplementary Fig. 5B).

### Role of caspases in Opto-Asc-induced apical extrusion

Opto-ASC efficiently induced cell death in periderm and basal cells, but not in muscle cells which the lack downstream effector caspases (Spead *et al*., 2018). To test the role of inflammatory caspases in speck-induced cell death, we disrupted the function of the two caspase-1 homologues, caspa and caspb, by outcrossing *Tg(Opto-asc)* to an existing caspa knock-out (KO) line (Kuri *et al*., 2017), but unexpectedly saw no effect on extrusion of periderm cells after Opto-Asc speck formation. We also used a CRISPR/Cas9 based method to create F0 knock-out (Kroll *et al*., 2021) as an alternative strategy of interfering with caspase expression The synthetic short guide RNAs (sgRNAs) targeting different exons of *caspa* and *caspb* (Fig. 7A) labeled in red in Fig. 7A were used in the double KO, and the ones labelled in grey for single gene KOs. The efficiency of the knock-out was validated by sequencing around all sgRNA binding sites.

**Figure 7:**
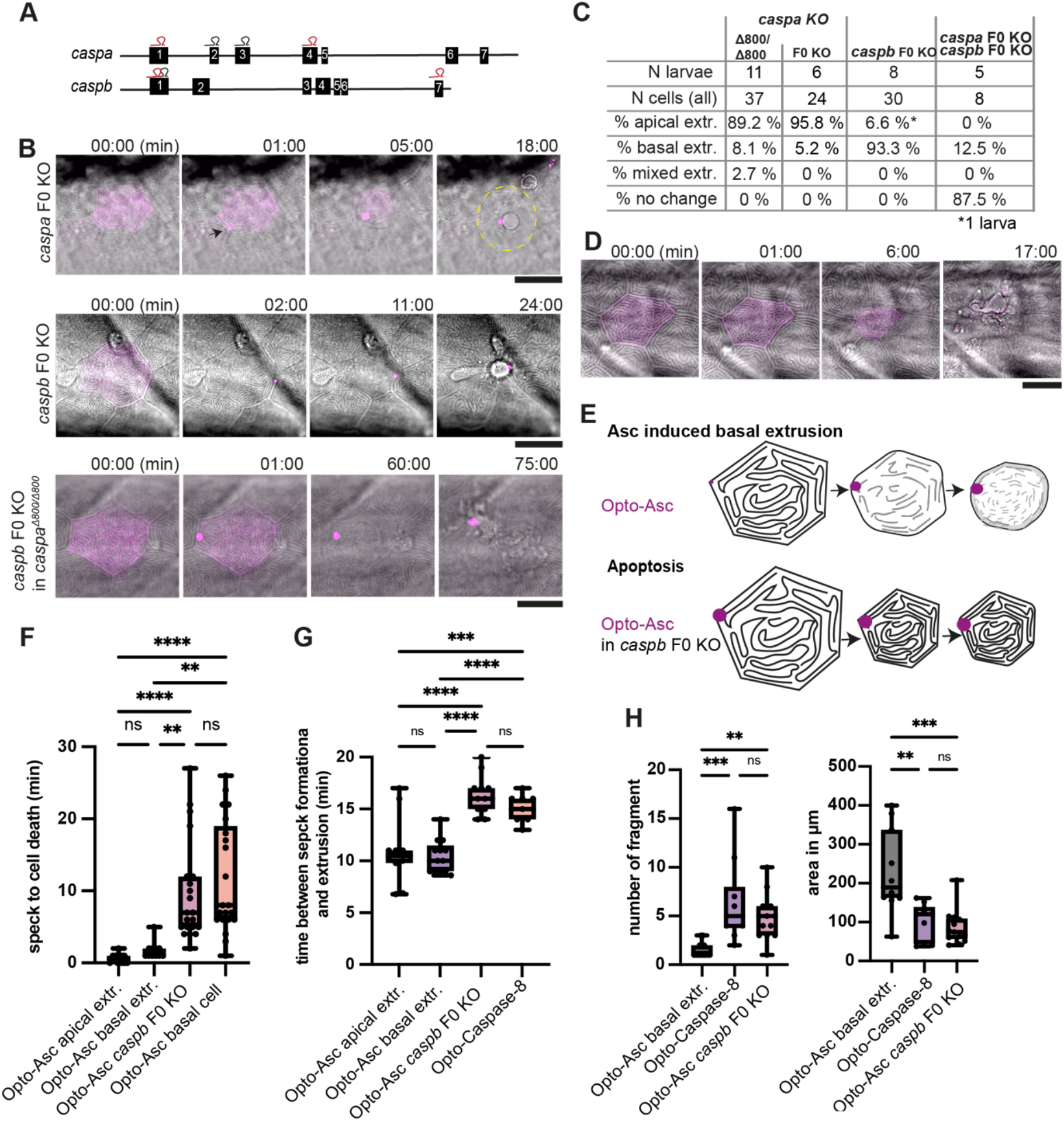
Inflammatory caspases in cell extrusion and rapid cell death after Asc-speck formation. **A**. Schematic of the *caspa* and *caspb* genes and sgRNA binding sites for F0 CRISPR/Cas9 knock-outs (F0 KO). **B**. Representative responses of periderm cells after speck formation in *caspa* F0 KO, *caspb* F0 KO and *caspa* F0 KO/*caspb* F0 KO. Scale bar is 20 µm. **C**. Percentages of apically extruded and basally extruded cells and cells that do not die within the observation time after Opto-Asc speck formation in different F0 KO backgrounds. **D**. Periderm cell dying by apoptosis after Opto-Caspase-8 induction; scale bar is 20 µm. **E**. Schematic of basally extruding cell after Opto-Asc induction and isomorphic shrinkage as seen in in *caspb* F0 KO larvae after Opto-Asc induction. **F**. Time between Opto-ASC speck formation and first morphological signs of cell death (deformation of cell) in the indicated conditions. **G**. Time of gap closure after death of single periderm cells which are either extruded or retained after the indicated treatments. **H**. Number of fragments at the time of gap closure and area of the largest cell fragment of retained cells after Asc-Speck formation in WT and *caspb* F0 KO larvae and of apoptotic cells after Opto-Caspase-8 induction. **F-H**.**=p<0.01, ***=p<0.001, ****=p<0.0001

Caspa deletion had no effect on the morphology or timing of Opto-Asc-induced cell death or the extrusion efficiency (around 90 % of cells were still apically extruded). However, periderm cells lacking Caspb were no longer extruded apically (Fig. 7B and C), but instead they displayed the apoptotic phenotype, similar to the one observed in these cells after Opto-Caspase-8 activation (Fig7D). In particular, these cells shrank while, retaining the pattern of actin ridges (Fig.7E).

Thus, either caspa or caspb alone is sufficient to trigger cell death, but only caspb can trigger apical extrusion.

In mammalian cells, inflammasome formation results in pyroptosis, which is defined as cell death induced by Gasdermin pore formation (Liu *et al*., 2016). The Zebrafish genome encodes two gasdermins (Gsdms): GsdmEa and GsdmEb, which can be cleaved in vitro by Caspb and apoptotic Caspase-8 and -3 respectively (Wang *et al*., 2020; Chen *et al*., 2021). To test the role of these Gsdms in Opto-Asc-induced cell death and extrusion, we used two ways of inactivating Gsdm function, a F0 CRISPR/Cas9 knock-out with four synthetic sgRNAs for each gene, which has shown to induced >95% protein-null alleles (Kroll *et al*., 2021) and the Gsdm inhibitor LDC7559 similar as was used in (Isles *et al*., 2021) to inhibit Gsdm pores of neutrophils. However, neither of these treatments had an effect on the phenotype or dynamics of cell death or on the polarity of extrusion (data not shown). We imaged and genotyped at min 5 larvae for single and the double knockout of Gsdms.

We also measured the time span between speck formation and first signs of cell death, in this case the start of cell deformation (shrinking) in different F0 KO backgrounds. F0 KO of *caspb* led to a delay in the response to speck formation of up to 20 min, regardless of the direction of extrusion (Fig. 7F). Double deletion of caspa and caspb delayed Opto-Asc-induced cell death up to 180 min or longer. Similar to cells in which apoptosis was induced by Opto-Caspase-8, these dying cells required a longer time (around 15 min) to be excluded from the periderm than during Opto-Asc induced cell death in control larvae (Fig. 7G).

We also observed differences in the number and size of cell fragments, as measured by the number and diameter of Akt-PH-GFP labeled membranes vesicles in basally extruded peridermal cellsdepending on whether death was induced by Opto-Asc with or without caspb or by Opto-Caspase-8 (Fig. 7H). Again, the phenotype of Opto-Asc-induced death in the absence of caspb resembled that of Opto-Caspase-8-induced apoptosis. We will therefore refer to this phenotype as opto-Asc-induced apoptosis.

### Ca^2+^ signaling in response to different types of cell death

Pyroptotic cells have been shown to release ATP and other signaling molecules in order to attract phagocytic cells (Wang *et al*., 2013), in order to see if this affects Ca^2+^ signaling in surrounding cells we looked at the Ca2+ signaling in the zebrafish skin in response to necrosic cell death and apoptosis. We used the Ca^2+^ signaling reporter GCamp6 to characterize the response of surrounding cells to cell death induced by Opto-Asc specks or Opto-Caspase-8. We crossed the *Tg(Opto-asc)* line to *Tg(ß-actin:GCamp6)*, a line expressing GCamp6 under the ubiquitously active ß-actin promoter(Chen, Xia, Michael R. Bruchas, *et al*., 2017) and subsequently imaged single cells expressing Opto-Asc and their immediate surrounding at high temporal resolution (>5 sec frame rate).

Without induction of cell death, only sporadic and weak Ca^2+^ signals were seen in individual cells at a frequency of about 0.2 per minute (3 cells in 15 min) within an imaging area of around 24 cells corresponding to 13 mm^2^ (Supplementary Fig.6A). In contrast, opto-ASC activation induced strong Ca^2+^ responses in neighbors. As an example, Fig. 8A shows Ca^2+^ signaling in periderm surrounding a cell being apically extruded following Opto-Asc speck formation. The speck-forming cell as well as the basal cell underlying the site of the speck showed increased Ca^2+^ levels within 15 sec of the speck becoming detectable; this preceded the shrinking of the cell, which started after around 30 seconds. Ca^2+^ signaling then spread to the surrounding cells in a wave-like manner. The first intense Ca^2+^ wave was immediately followed by a smaller second wave, after which the signal became sporadic over the course of 5 minutes and then subsided. Once the dying cell started blebbing, its internal Ca^2+^ level remained high until lysis, indicating the loss of its ability to control intracellular Ca^2+^ homeostasis.

**Figure 8:**
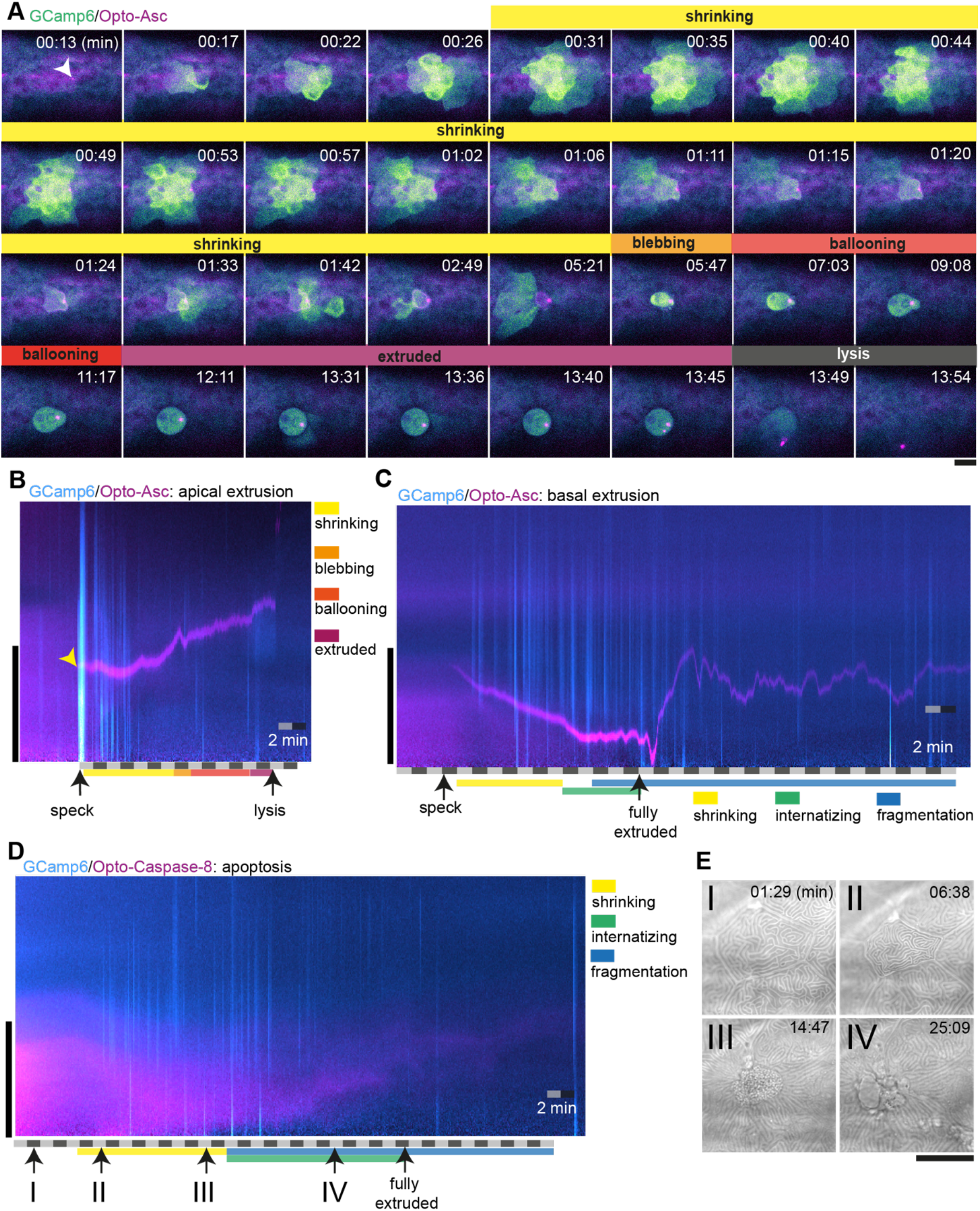
Ca^2+^ signaling in epithelial cells in response to dying cells. **A**. Example of Ca^2+^ signaling in epithelial cells surrounding a cell forming an Opto-Asc speck (yellow arrow head) and being extruded from the tissue. Frames from a time-lapse image sequence of a larva expressing GCamp6 (light green) and Opto-Asc (magenta). Phases of the extrusion were characterized using the bright field channel, as described in Fig 5 and are color coded (yellow-black). Scale bar is 20 µm. **B-D**. Two-dimensional representation in time and space (derived from 3D kymographs) of the Ca^2+^ response to cell death in the larval epidermis. The y-axis shows the radial space around the dying cell, as described in supplementary figure 7. GCamp6 Ca^2+^ sensor signal intensity is shown in cyan and Opto-Asc in magenta. The scale bar on the y-axis is 20 µm. Time is marked along the x-axis in 2min blocks. **B**. Ca^2+^ signaling response to Opto-Asc-induced apical extrusion as shown in A. The phases of extrusion are color coded (yellow to dark red) as in panel A and Figure 5C. **C**. Opto-Asc-induced cell death with basal extrusion. Phases of extrusion marked along the x-axis. **D**. Opto-Caspase-8 (magenta)-induced apoptosis. Numbers on the x-axis refers to the images in E. **E**. Bright field images of the cell analyzed in D illustrating stages of apoptotic cell death. I: Cell morphology before the first signs of cell death; II: isomorphic shrinkage of cell; III: microridge pattern has dissolved and cell starts to internalize; IV: cell has started to fragment. Scale bar in all images is 20 µm.

To analyze these observations in a more quantitative manner, we used a two-dimensional representation of time and space (derived from 3D kymographs; see Supplementary Fig. 7) to represent the calcium response in the area around the dying cell (Fig. 8B). The sum of the signals along a radial sweep around the cell is represented on the y-axis (with the center of the dying cell at y = 0) versus time on the x-axis (Supplementary Fig. 6B-D). The initial Ca^2+^ waves (in cyan) appear as strong lines following the appearance of the speck (yellow arrow), and the sporadic Ca^2+^ signals of surrounding cells appear as the weaker lines. The speck is visible following its appearance as a magenta line, until it disappears following cell lysis. The extruded cell appears as a cyan shadow once extruded, and remains visible until lysis. Cells that were basally extruded responded to speck formation with immediate increased Ca^2+^ levels in 3 out of 5 cases. For the first 15 minutes after the cell started to shrinkwe detected on average 8.6 cases of increased Ca^2+^ signaling in basal cells and 30.7 in periderm cells (N=3 cells of different larvae) (Supplementary Fig. 7A and B).

By contrast, caspase-8-induced apoptosis of periderm cells did not induce a comparable Ca^2+^ wave in surrounding periderm (Fig. 8D), and apoptotic cells themselves never showed increased Ca^2+^ levels. During the shrinking phase, before the cell was fully internalized, the surrounding cells (mainly basal cells) showed only sporadic Ca^2+^ signaling. We detected on average 21 cases of increased Ca^2+^ signaling in basal cells and 12.3 cases in periderm cells directly adjacent to the dying cell within the first 15 minutes after the cell started to shrink (N=3 cells of different larvae) (Supplementary Fig. 7C). Finally, when periderm cells died via Asc-induced apoptosis in Caspb knock-outs, the Ca^2+^ response of surrounding cells and the dying cell itself was similar to the response to Opto-Caspase-8-induction in the periderm. In summary, the response of the surrounding cells depended on the direction of extrusion and on the stimulus that triggered cell death. A strong calcium response in the form of a wave in neighboring periderm cells and underlying basal cells occurred only in the case of ASC-induced extrusion and most prominently when cells were apically extruded.

## Discussion

Our results demonstrate that opto-ASc is an efficient tool to induce inflammasome formation and ASC-dependent cell death in zebrafish. Compared to Asc over-expression used in our earlier studies (Kuri *et al*., 2017), or I infection models (Forn-Cuní, Meijer and Varela, 2019), it allows a more precise spatial and temporal manipulation of inflammasome activation and cell death. Heat-shock induced expression of Opto-Asc is highly variable between cells within individual larvae and between larvae, which has both advantages and disadvantages. Mosaically distinct Asc levels allow the assessment of the role of Asc levels and the response of neighboring cells within the same experimental animal.

If a more uniform expression of Opto-Asc were desired, this could be achieved by the expression of Opto-Asc under the control of tissue-specific promoters combined with Cre-Lox or trans-activator-induced expression (Knopf *et al*., 2010; Mosimann *et al*., 2011; Gerety *et al*., 2013).

Opto-Asc oligomerization is efficiently induced within minutes by constant exposure to 488 nm light. After a single light pulse, specks were often not observed immediately, but appeared with a delay of up to 40 min. This is unexpected, because it is unlikely that Cry2-olig would remain active for so long that dimerization could be delayed for more than a few minutes (Taslimi *et al*., 2014), and one would expect that even a single initial dimerization of Opto-Asc would have led to the immediate recruitment of the endogenous Asc that is available throughout the cell. Our observations may suggest that the initial seeding event could have created a dimer that was not yet active, but that a second step of maturation could subsequently occur. This highlights the fact that we still know very little about the unusual dynamics of speck assembly, or the mystery why the cell normally always forms only one single speck.

In response to the formation of an Opto-Asc speck periderm cells died in two morphologically different ways. Cells were extruded either apically or basally, and some cells were simultaneously extruded in both ways. Apical extrusion of periderm cells has been described not only in response to infection, but also in response to treatment with the antibiotic geneticin (G418) (Eisenhoffer and Rosenblatt, 2011) and during tissue homeostasis as a response to cell crowding (Eisenhoffer *et al*., 2012). Apoptotic stimuli like geneticin have been shown to induce extrusion via sphingosine-1 phosphate (S1P) and its receptor, which is expressed in the surrounding cells. We have previously shown that S1P signaling does not seem to drive extrusion of pyroptotic cells in Caco-2 cell co-cultures (Shkarina *et al*., 2022), but so far it is unclear if S1P signaling or another mechanism activated downstream of the inflammasome is responsible for apical extrusion. Although the factors determining the direction of extrusion remain to be discovered, the different ratios of apical and basal extrusion of periderm cells in different zebrafish strains suggest a genetic component.

These findings indicate that different types of cell death are not entirely separable, but that they represent different manifestations of parallel and interconnected signaling pathways, the overall balance of which defines the final outcomes. This is also evident from other studies in the past, where disrupting one pathway directs the cell towards a different pathway that may have a different phenotypic outcome (Tsuchiya *et al*., 2019). Perhaps it is only through experimental activation of the most downstream effector that a ‘pure’ phenotype can be induced.

In the case of zebrafish periderm, we identified Caspb as essential for apical extrusion. In the absence of caspb, cells die with an apoptotic morphology in response to speck formation. The requirement for caspb for apical extrusion and lysis might also explain the difference in the response to Asc-speck formation between the periderm and the underlying basal cells. According to cell type specific RNA sequencing data (Cokus *et al*., 2019), basal cells express ten times lower levels of Caspb than periderm cells.

Although Caspa localizes to the Asc speck (Kuri *et al*., 2017) and is necessary for rapid cell death if no Caspb is present, we did not find it to be necessary for speck induced extrusion. This contradicts our own previous work, in which we identified caspa as responsible for fast extrusion after speck formation (Kuri *et al*., 2017). We have now re-analyzed our previous data (Supplementary Fig 8) and found that the delay between speck formation and cell death can be partially explained by the type of cell that was imaged, namely basal cells although we could observe a much stronger delay between speck formation and cell death compared to WT basal cells.

Zebrafish Caspa and Caspb are thus redundant in their role as rapid cell death inducers, but the downstream mechanisms that lead to either extrusion or apoptosis of periderm cells are yet to be identified.

Apoptosis and Asc-induced cell death are both initiated within a short time after speck formation and cause the recruitment of actin in surrounding cells to the side facing the dying cells. The closure of the surrounding epithelium above or below the extruded cell is faster for Asc-induced extrusion than for apoptosis. This difference in rate of wound closure could be caused by different dynamics in the cell itself, or in the surrounding cells. The fact that we observe an immediate Ca^2+^ response in the case of Asc-induced death, but not apoptosis, indicates signalling from the dying cell to the surrounding epithelium, and this signal differs between cells dying apoptotically or by Asc-induced death The dependence on the fast apical extrusion and the rapid Ca^2+^ response require Caspb, pointing to a Caspb-dependent signalling event. A similar reaction was observed in cultured intestinal epithelial cells in which a bacterial infection sensed by the NAIP/NLRC4 inflammasome induce a fast myosin-dependent contraction in surrounding cells. This reaction was dependent on the activation of sub-lytic GSDMD pores and the resulting ion flux and independent of the cell death or extrusion of the cell (Ventayol et al., 2021).

Zebrafish do not have a direct homolog of GsdmD but two homologues of GsdmE (Gsdmea/b). Human GsdmE was shown to be activated downstream of caspase-3 during apoptosis (Wang *et al*., 2017) and it plays a role in the caspase-3 mediated lytic death of primary human keratinocytes after viral inactivation of the Bcl-2 pathway (Orzalli *et al*., 2021). Both zebrafish gasdermins can be cleaved by caspb and other apoptotic caspases (but not caspa) *in vitro* (Chen *et al*., 2021). It was therefore surprising to find that we did not see a change in peridermal keratinocyte cell death after ASC-speck formation when we knocked out Gsdme a and b by CRISPR/Cas9, or treated fish with a Gasdermin inhibitor. Although our CRISPR/Cas 9 strategy is highly efficient for other genes, we cannot rule out that Gsdme a and b are redundant in their function to mediate Caspb dependent extrusion and that a relatively small amount is sufficient to induce this form of cell death. Additionally, we find that cells which are extruded do not lyse immediately upon swelling which is normally the case in pyroptotic cultured cells (Shkarina *et al*., 2022). Since we find a caspb-dependent difference in cell death phenotypes, the question arises which other component could cause the apical extrusion of cells and wave-like Ca^2+^ signaling response in surrounding cells. Possible candidates could be pannexins, especially pannexin1a which is expressed in skin cells (Cokus *et al*., 2019) and is necessary downstream of Caspase-11 to induced pyroptosis in response to LPS in mouse macrophages (Yang *et al*., 2015). The release of ATP caused by pannexin pores might explain the strong calcium wave in caspb-dependent extrusion of peridermal keratinocytes (Mori *et al*., 2022).

## Materials and Methods

### Zebrafish husbandry, transgenic lines, and genotyping

Zebrafish (*Danio rerio*) were cared for using standard procedures as described previously (Westerfield, 2000) and in accordance with EMBL guidelines. All experimental procedures were approved by the EMBL Institutional Animal Care and Use Committee (IACUC nos. 2019-03-19ML). We used the following zebrafish wildtype lab strains: AB2B2, AB, golden and Wild Indian Karyotype (WIK). The *Tg(HSE:mCherry-Cry2olig-asc)* line was generated by co-injecting embryos at the one-cell stage with Opto-caspase plasmids with transposase mRNA (100 ng/μl). To follow the actin dynamics in skin cells, we used the *Tg(krt4:Gal4) line* (Wada *et al*., 2013) crossed to *Tg(6xUAS:mNeonGreen-UtrCH_6xUAS:lyn-tagRFP)* generated in the lab of D.Gilmour by Jonas Hartmann. Actin dynamics in basal skin cells were visualized in the *Tg(rcn3:Gal4*) line (Ellis, Bagwell and Bagnat, 2013) crossed to *Tg(6xUAS:mNeonGreen-UtrCH_6xUAS:lyn-tagRFP)*. To induce specks and follow endogenous ASC we used *Tg(HSE:asc-mKate2)* and the endogenously tagged asc:asc-GFP line (Kuri *et al*., 2017). To induce apoptotic cell death, we used *Tg(HSE:mCherry-Cry2olig-caspase-8)* (Shkarina *et al*., 2022). For monitoring Ca^2+^ signaling we used *Tg(ß-actin:GCamp6)* (Chen, Xia, Michael R Bruchas, *et al*., 2017).

### Cloning of Opto-Asc and Opto-Nlr variant constructs and transient expression

The Cry2olig-mCherry construct (plasmid #60032) was obtained from Addgene, and plasmids containing Asc, Asc(4xmut), Pyd, Nlr and Card published in (Kuri *et al*., 2017) were used as cloning templates. The initial HSE:mCherry-Cy2olig ASC construct was cloned as described in (Shkarina *et al*., 2022). Other constructs containing ASC variants (PYD and CARD) used the HSE:mCherry-Cy2olig-Asc as background and were modified by seamless cloning using the NEB Assembly Kit, after SmaI/EcoRV or XmaI/EcoRV digestion of the plasmid (replacing Asc). The following constructs were generated and used for transient expression: HSE:mCherry-Cry2oligR489E, HSE:mCherry-Cry2olig-Pyd, HSE:mCherry-Cry2olig-Card, HSE:mCherry-Cry2olig-Nlr-FL, HSE:mCherry-Cry2olig-Nlr-Pyd. All inserts were verified by sequencing, and constructs are deposited in the European Plasmid Repository (https://www.plasmids.eu)For expression of transient constructs, we co-injected expression plasmids with Tol2 transposase mRNA (100 ng/µl) in embryos at the one-cell stage.

Larvae were screened at 2.5 dpf for expression of tag:RFP in the heart as described in (Kuri *et al*., 2017); For heat-shock driven expression, we incubated larvae at 2.5-3 dpf in 2 ml tubes in a heating block at 39°C.

### CRISPR/Cas9-mediated asc zebrafish knockout

We deisgned a custom gBLOCK (Integrated DNA Technologies) incorporating a guide RNA-targeting sequence preceded by a T7 promoter sequence. Guide RNAs against exon 1 of *asc* were used with the following sequences: 5′-GGTGGAGATCGAAGATCAAG-3′ and 5′-GCAGCTGCAGGAGGCTTTTG-3′ respectively. Guide RNAs were synthesized using MEGA shortscriptTM Kit (Applied Biosystems), according to the manufacturer’s protocol, and were purified using RNeasy Mini Kit (Qiagen, #74106). Cas9 mRNA was synthesized using mMESSAGE mMACHINE® SP6 Transcription Kit (Thermo Fisher Scientific, #AM1340). 2 nL of a mixture containing 250 ng/μL gRNA and 0.1 mg/mL Cas9 protein was injected into the yolk of 1-cell AB zebrafish embryos.

Genotyping was performed by extracting genomic DNA from fin clips or larvae using the QuickExtract DNA extraction solution (Epicentre) and amplifying loci using EMBL in-house Phusion polymerase, primers are listed in Supplementary Table 1.

### F0 CRISPR Screen

The F0 CRISPR screen was performed as described in (Kroll *et al*., 2021) for caspa, caspb, *gsdmea, gsdmeb* and *caspase-8a*. sgRNA target sites were selected for high predicted cleavage efficiency within various exons of candidate genes using CCTop (Stemmer *et al*., 2015).

Sequences of synthetic sgRNAs and gene identifiers are listed in Supplementary Table 1. We used 3-4 synthetic sgRNAs (Sigma) per gene, which were diluted in water to a concentration of 100 µM. Cas9 protein was produced by the EMBL PEPCore facility and stored in a concentration of 62,5 µM in Cas9 buffer (20 mM Tris-HCl, 500 mM KCl and 20 % vol glycerol). For each experiment, we freshly mixed equal volumes of sgRNA and protein solution (1 µl sgRNA per 1 µl Cas9) and injected the mixture into early embryos within the first 15 min after fertilization. All CRISPR KOs were confirmed by PCR and sequencing around target sites.

### Live imaging and optogenetics

Zebrafish larvae were anaesthetized using 4mg/ml tricaine pH 7 and mounted on their right side in 0.75% low melting agarose on MaTec glass bottom petri dishes (35mmNo. 1.5 coverglass, 0.16-0.19 mm). Live imaging was performed at 20°C using a Zeiss LSM780 NLO confocal microscope (Carl Zeiss) and a 40× C-Apochromat (NA 1.20) water immersion objective (Carl Zeiss). To image GFP, mNeonGreen and the GCamp6 Ca^2+^ reporter, we used the 488 nm multiline Argon laser (Lasos Laser GmbH). To image mCherry and lyn-tagRFP, we used a HeNe laser at 561 nm. For detection of the far-red dye DRAQ7 we used the HeNe 681 laser.

We determined the laser intensity for the 488 nm laser at regular intervals using a chroma slide at a fixed laser power of 5 %. The total laser power was measured using a laser power meter (THORLABS GmbH Dachau Germany SN: P0023941) equipped with the sensor S121C 400-1100nm with a capability of measuring 500mW. Whenever not stated otherwise, we used 5 % 488 argon laser power (24.8microWatt/pixel) to stimulate oligomerization of optogenetic constructs.

For 2-photon activation of Opto-ASC in single cells, we used a 140-fs pulsed multi photon laser (chameleon; Coherent) of the LSM780 NLO and operated the microscope using the Zen Black Software (Carl Zeiss, 2012 version) together with the Pipeline Constructor Macro (Politi et al., 2018).

For overnight imaging of whole larvae, we used a 20x air objective (Carl Zeiss) and acquired multiple stacks (2×7) to cover the entire larva. Stacks were then stitched together using the Zen Black software. Stacks were acquired every 15 minutes for 12-16 hrs.

For high-resolution time-lapse imaging of the Opto-ASC speck, Opto-PYD and Opto-CARD we used a Zeiss 880 Airyscan and imaged in Airy fast mode with a C-apochromat 40x/1.2W Korr FCS M27 objective. For activation of opto-constructs and detection of GFP we used 0.2% 488nm and for detection of mCherry 1.5% 561nm. To detect the GCamp6 signal, we optimized imaging conditions to reach a frame rate below 5 sec. The stack size was 8 µm (1 µm z resolution).

### Image analysis and quantification

We used Fiji ImageJ for image analysis (Schindelin *et al*., 2012). Images acquired on the Zeiss 880 Airyscan were processed with Zen black software (Zeiss). We used the cell counter plugin to manually count total numbers of periderm cells and numbers of periderm cells expressing Opto-Asc. Example images are always presented as maximum intensity z-projections. The total mean fluorescence of larvae was measured in maximum z-projections in the region of interest (ROI) manually defined by the outline of the larva. To determine the expression of Opto-Asc in single cells, we chose the three z-planes in which most of the cell was visible and measured MFI in a selected region with uniform expression in an average intensity z-projection.

Line profiles for high resolution images of specks were normalized by dividing by the average gray value and subtraction of the lowest gray value. For measuring speck size, a line was drawn through the longest diameter of the speck; the diameter was determined by measuring the distance between the two points of maximum slope of the plot profile along the line.

To determine the growth of specks, we marked the outlines of the speck in each time frame for Opto-Asc and asc-GFP separately and measured the area. We started the measurements at the first time point where mCherry/GFP visibly aggregated. We calculated the time difference starting from the appearance of the first mCherry/mKate2 aggregates to the time point where total intensity (intensity * area) of the speck reached a constant value.

To determine if a periderm cell was extruded apical, basal or in both directions we used the bright field imaging, anything that is retained below the periderm was consider as basally extruded, everything that left the tissue and dissociated from the epithelial layer to the water was considered apically extruded. In case of mixed extrusion part of the cell was retained the other part apically extruded.

For estimating the time between speck formation and initiation of cell death, we calculated the interval from first detectable Opto-Asc aggregates to the time when the cell outline first deformed. For the analysis of Ca^2+^ signaling of bystander cells in response to cell death, we generated Kymographs for a circular area around the dying cells using the Radial Reslice Fiji plugin (Cooper, 2009) which measures the pixel intensity along a line rotating around a center in 360 degree. By creating an average projection of all lines, we created a two-dimensional representation of the area around the cell for every time point. The exact procedure with examples images is shown in Supplementary Fig. 6.

To quantify the Ca^2+^ response of bystander cells to basal extrusion and apoptosis, we manually counted the number of periderm cells and basal cells which showed a GCamp6 signal for every minute after initiation of cell death for 15 min.

### Statistics

All statistical analysis was done using Prism (version 9) (Graphpad). P-value ranges are included in figures as follows: * is p<0.05, ** is p<0.01 *** is p<0.001 and **** is p <0.0001. For comparison of different heat shock conditions (Fig. 2B and C) and the comparisons in Fig. 7F-H, we used ordinary one-way ANOVA. For determining correlations, we used simple linear regression. To compare average MFI between speck forming and non-speck forming cells, we used a Wilkoxon Test. To compare sizes of specks and time of speck formation, we used a Mann-Wittney test. For the difference in time shown in Fig. 4C and Fig. 5E, we assumed unequal standard deviations and used a t-test with Welch’s correction.

### Chemical treatments

For the detection of cell lysis, we used Draq7 (Invitrogen), a far-red dye that only stains nuclear DNA once the membrane integrity is lost. We preincubated fish in 0.3 µM (1:1000) Draq7 in E3 fish medium for 1 hr and added the dye directly to the low-melting agarose and the fish medium in which larvae were imaged, to a concentration of 0.3. µM.

To inhibit gasdermin pore formation, we treated 2dpf larvae over night with 10 µM to 100 µM Gasdermin D inhibitor LDC7559 (MedChemExpress). The inhibitor was added directly to the fish water as described in (Isles *et al*., 2021), and larva were imaged at 3 dpf.

## Acknowledgements

The authors thank the Advanced Light Microscopy Facility (AMLF) at the EMBL-Heidelberg for their continued support and Manuel Gunkel for his help with laser intensity measurements. The authors thank Darren Gilmour and Jonas Hartmann for providing the *Tg(6xUAS:mneonGreen-UtrCH)* zebrafish line. We thank Alexandre Paix for providing guidance and material for CRISPR/Cas9 experiments and Takehito Tomita for help with analysis of GCamp6 imaging data.

## Author’s contribution

Author contributions: EH and ML conceptualized the study. KS and PB designed and created initial optogenetic tools. EH designed in vivo experiments. EH and SD performed in vivo experiments. BR and JRX created the Asc (Pycard) knockout fishline. All authors contributed to data analysis and manuscript writing.

