## supplementary figures for "The Opto-inflammasome in zebrafish: a tool to study cell and tissue responses to speck formation and cell death"

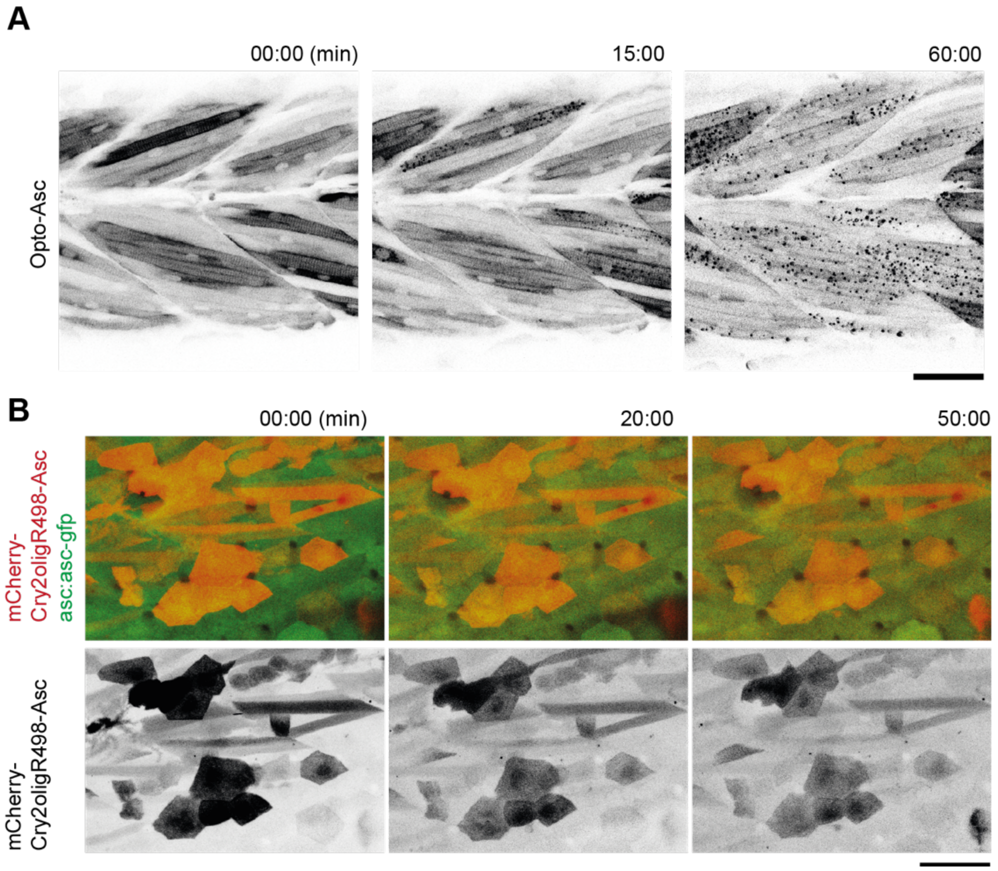


***Supplementary figure 1: Control experiments for speck formation in muscle cells with Opto-Asc and with mutated Cry2olig in periderm cells.***

**A.** Opto-Asc*Tg(mCherry-Cry2olig-asc)* specks induced by blue light (488nm) in muscle cells in 3dpf larva. Multiple oligomers are formed by the end of the movie (60 mins). Opto-Asc is in black. Scale bar is 50 μm.

**B.** Periderm cells in a 3dpf larva expressing mCherry-Cry2oligR498-Asc (in red). The mutated version of Cry2olig does not form oligomers even after 50 minutes of constant illumination by blue light (488nm). Endogenous Asc-GFP in green. Scale bar is 50 μm.

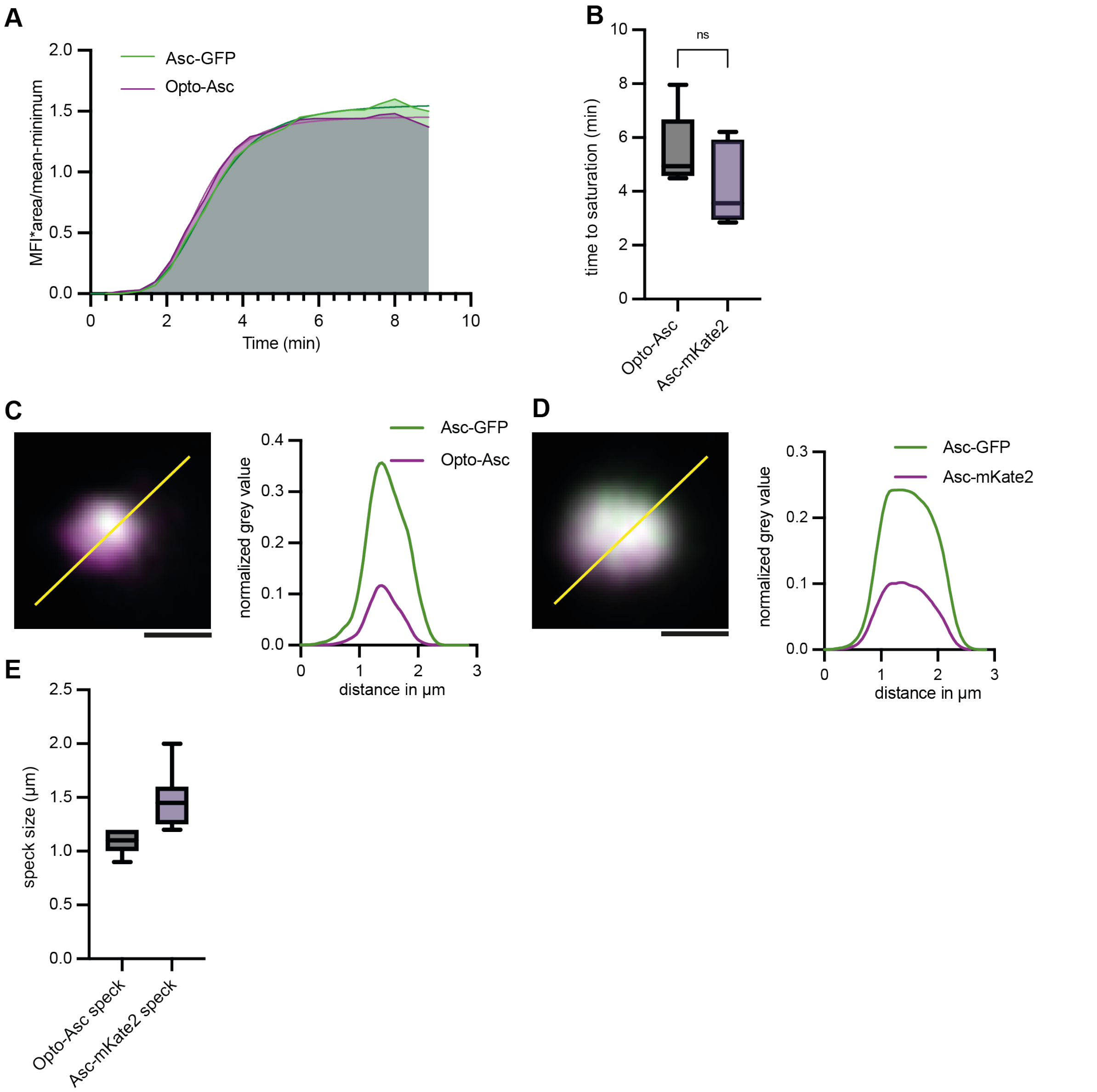


***Supplementary figure 2: Recruitment dynamics in Opto-Asc specks and Asc-mKate2 specks.***

**A.** Recruitment dynamics of endogenous Asc (Asc-GFP) and Opto-Asc to a speck induced by blue light (488nm). MFI was measured for GFP and mCherry at each time point over the entire area of the growing specks. Representative curve for 10 specks measured in 3 different larvae.

**B.** Recruitment of Asc-mKate2 and Opto-Asc in Asc-mKate2 specks and Opto-Asc specks. MFI of mKate2 and mCherry was measured at each point over the entire area of the growing speck and the time point at which the MFI value was constant was recorded.

**C.** Overlap of Opto-Asc (magenta) and endogenous Asc-GFP in light-induced speck at high resolution (Airyscan image acquired by Zeiss LSM 880).

**D.** Overlap of Asc-mKate2 (magenta) speck with endogenous Asc (GFP). The line ROI is drawn across the two specks and the normalized grey values are plotted along the line ROI. The distribution of the endogenously tagged Opto-Asc with Asc-GFP is uniform similar to that of the speck of Asc-mKate2 and Asc-GFP.

**E.** Comparison of diameter (in µm) of Opto-Asc specks (n=11) and Asc-mKate2 specks (n=8)

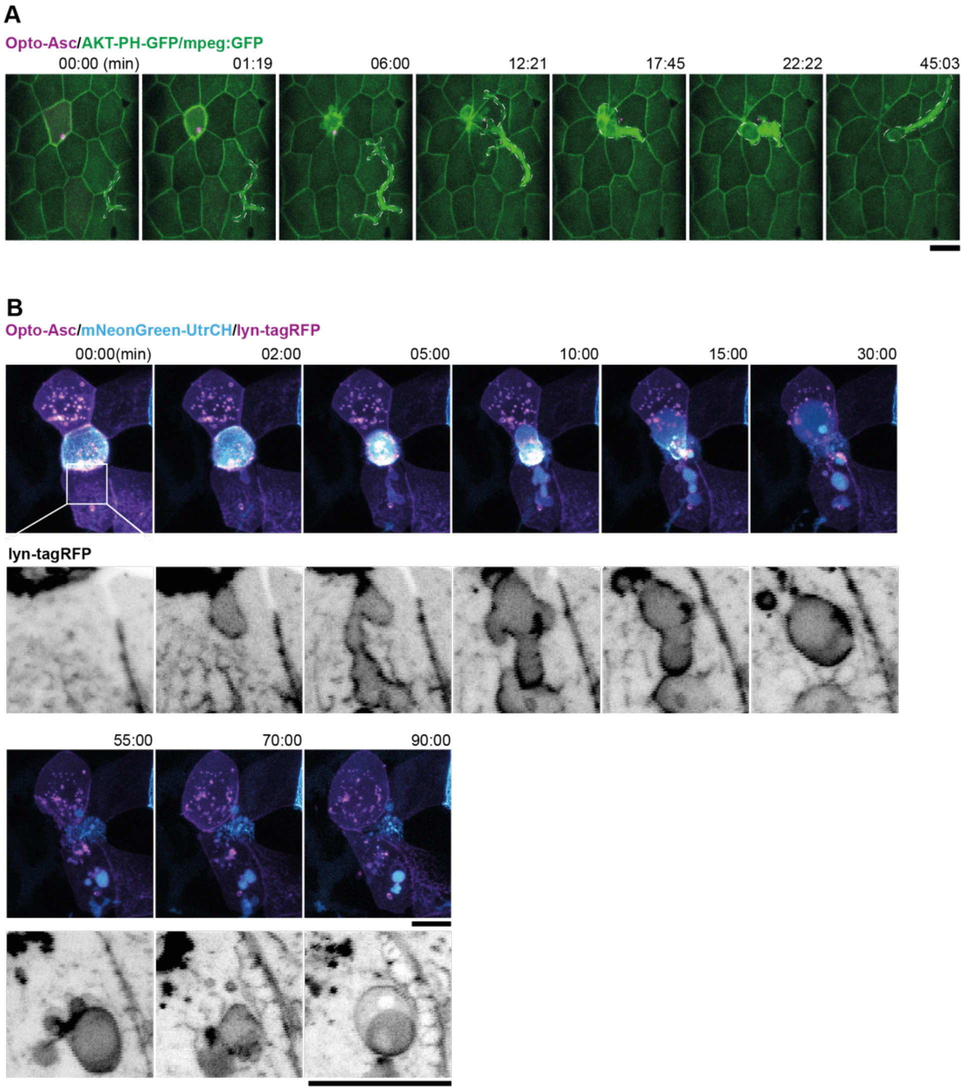


***Supplementary figure 3: Response of neighbouring cells to cell death after Asc-speck formation***

**A.** Response of macrophage to a basally extruded periderm cell (Opto-Asc in magenta). The macrophage (marked in yellow dashed outline) moves towards the dying cell which starts shrinking at 1:19 min, and engulfs the cell completely by 22:22 min. The macrophage is labelled with mpeg-gfp, and the periderm cell membranes with AKT-PH-GFP (both green). The scale bar is 20 µm.

**B.** Response of the surrounding periderm to a basally extruding periderm cell after Opto-Asc (magenta) speck formation. The white box marks the region of contact between the dying cell and a neighboring periderm cells. The same area is shown at higher magnification below with only the membrane label in grey scale. Blebs start to form as the dying cell disintegrates, and the remaining cell debris appears to be taken up by the surrounding cells. Actin is labelled by mNeonGreen-UtrCH (cyan) and the cell membranes by lyn-tagRFP (magenta).

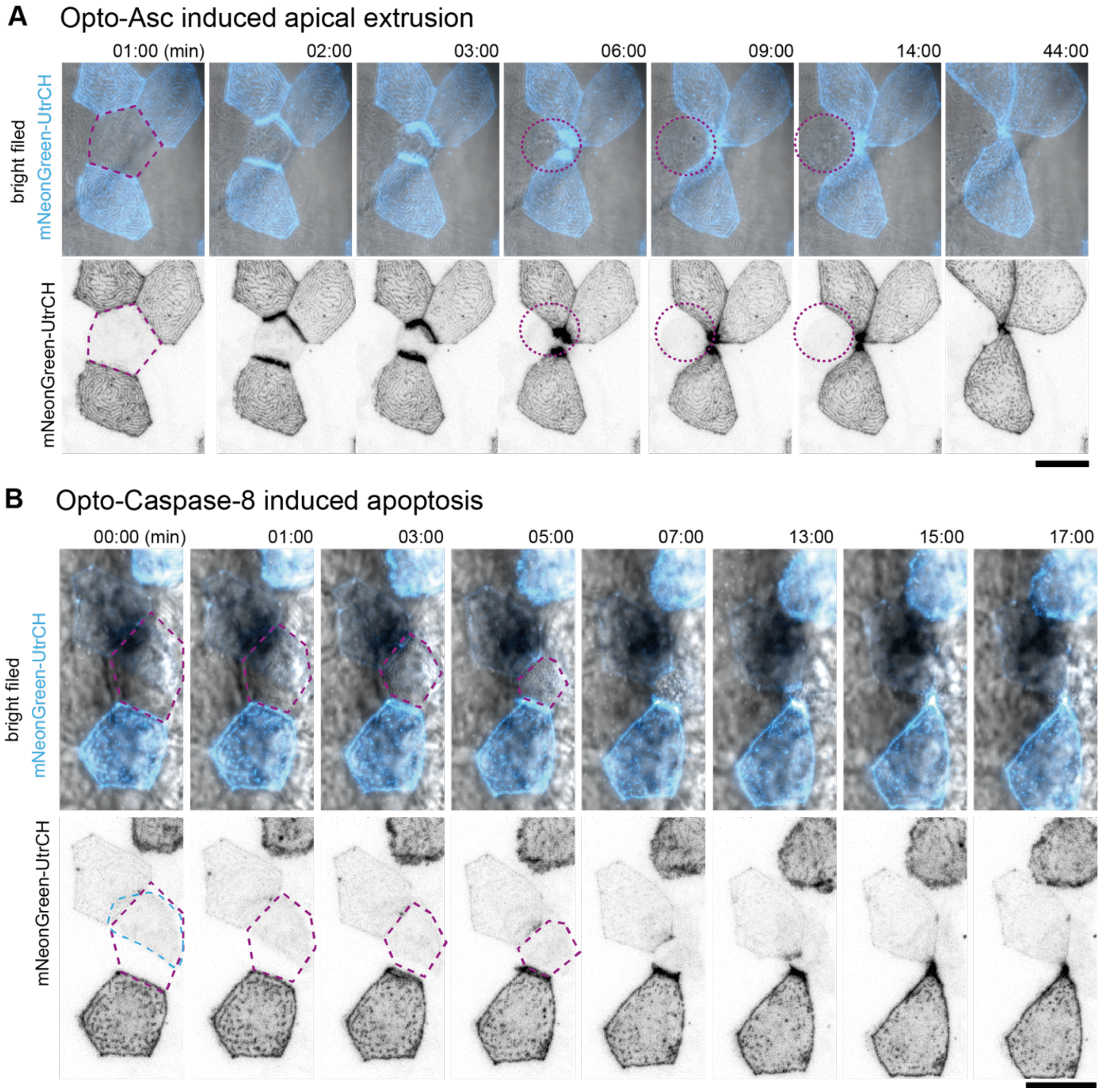


***Supplementary figure 4: Actin rearrangement in periderm cells near dying cells.***

**A, B:** Actin, labelled by mNeonGreen-Utr-CH (cyan in top row and black in the bottom row) is expressed in a mosaic pattern. As the surrounding cells close the gap over the dying cells, actin accumulates at the lateral membranes which are in contact with the dying cell.

**A.** Response of neighboring periderm cells to Opto-Asc induced apical extrusion. Opto-Asc is induced in the periderm cell marked in purple dotted line.

**B.** Response of neighboring cells to apoptosis induced by Opto-Caspase 8 in the periderm cell marked in purple. This cell does not express mNeonGreen-Utr-CH, but the signal comes from an underlying basal cell (outlined in blue).

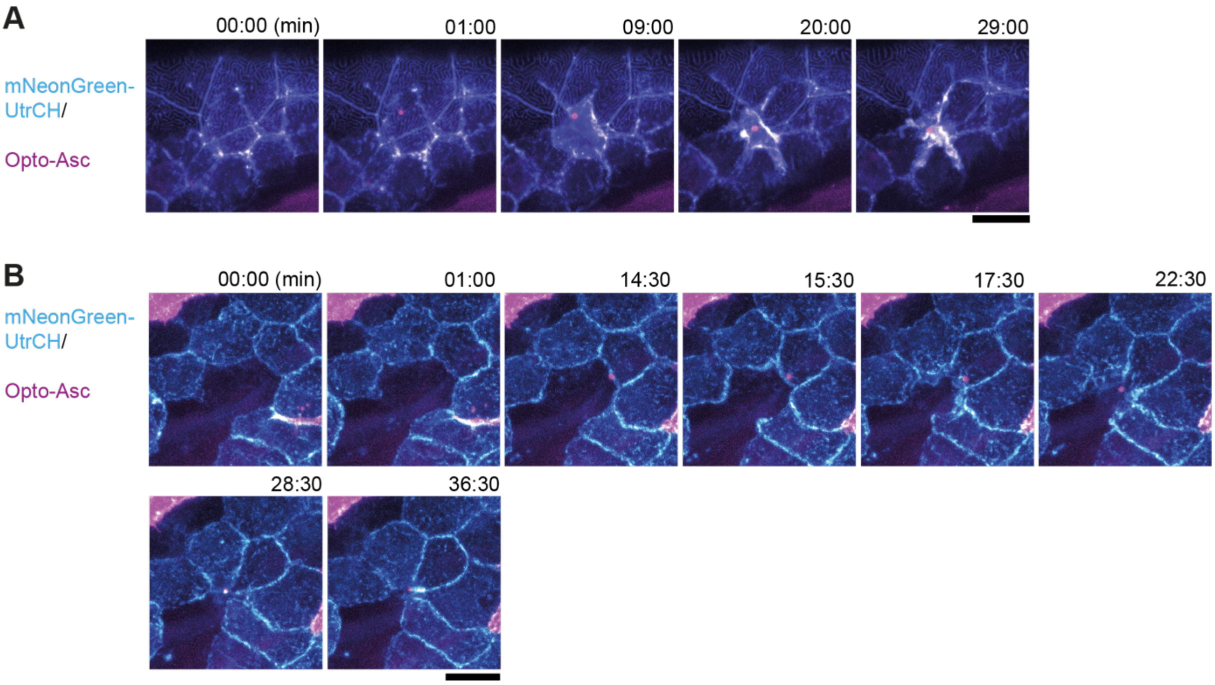


***Supplementary figure 5: Actin dynamics in neighboring cells after Opto-Asc-induced cell death in basal cells.***

**A, B:** *Tg(Opto-asc)* (magenta) was crossed with *Tg(UAS:mNG-UtrCH)* to label actin (shown in cyan).

**A.** Opto-Asc is induced in a basal cell. The speck is seen within 1 min. Actin concentrates at the periphery of the dying cell, but it is not clear whether it is in the dying or the neighboring cells. No actin rearrangement is observed in the periderm above the dying basal cell.

**B.** An Opto-Asc speck is induced in the central basal cell which does not express UAS:mNG-UtrCH. The speck is first seen at 14:30 minutes. The surrounding cells form lamellipodia that fill in the gap above the dying cell, but no strong actin concentration is seen, indicating that the actin at the periphery of the cell in panel A is the actin of the dying cell. Scale bar: 20 μm

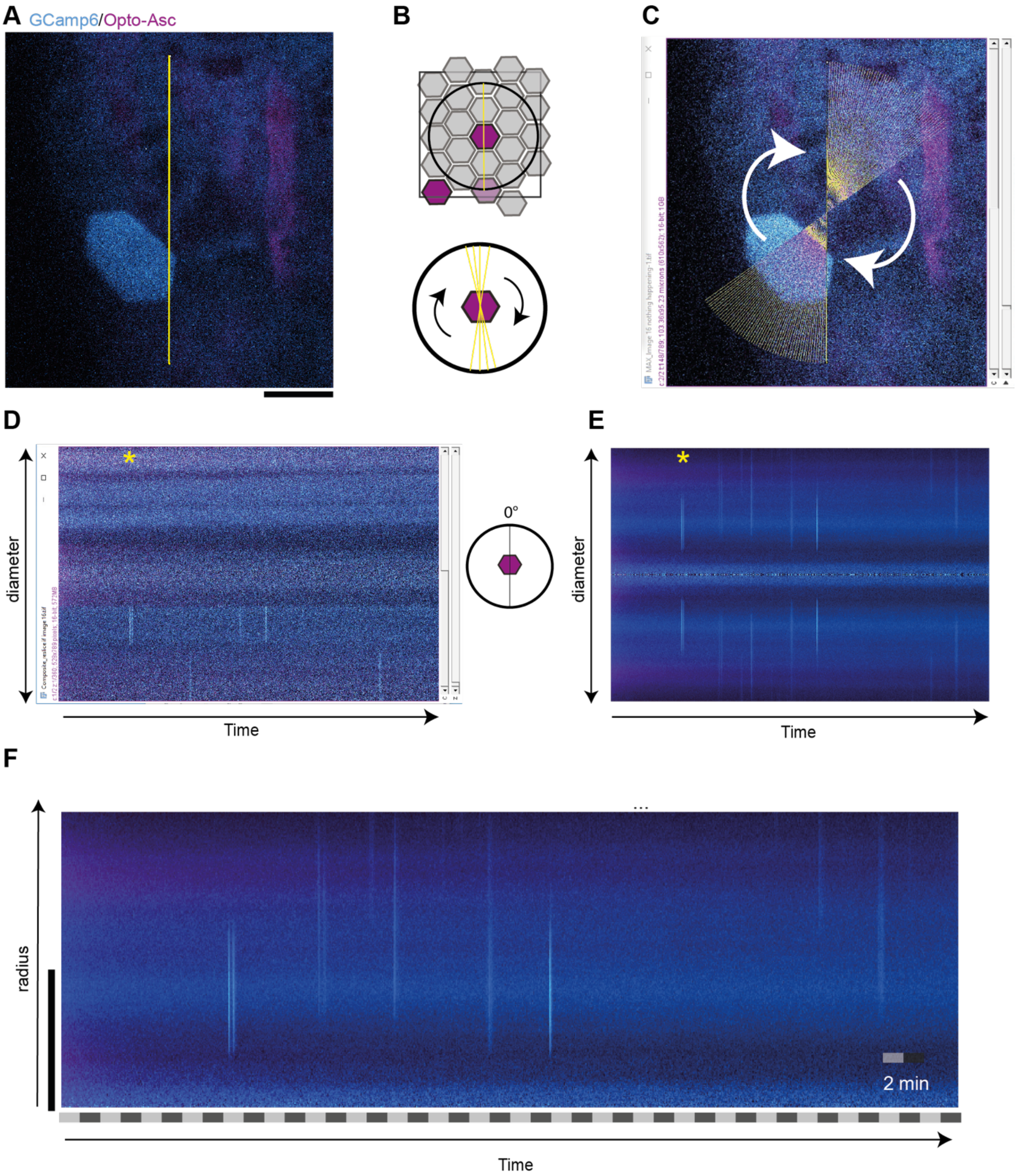


***Supplementary figure 6: Method for scoring calcium signalling in time and space and control for background signalling.***

**A to F:** Analysis of control larva expressing β-actin:GCamps (cyan) and Opto-Asc (magenta), yellow line: ROI; Scale bar in all panels: 20 μm

**A.** Imaged region with the cell of interest (which is not activated in this control) is in the center. The yellow line shows the line ROI at 0° along which fluorescence intensity is measured.

**B.** Top: Cartoon of the periderm. The circle around the cell of interest outlines the area that is analyzed in the kymograph.

Bottom: A line ROI is drawn across the center of the cell. This line is swept by 360 degrees in 1° degree steps to cover a 82.19 μm diameter circle surrounding the center of the cell.

**C.** Screen shot taken during the experimental course of measurement (at the 56° measurement). The fluorescence intensity at each point along the diameter is measured at each angle.
**D.** Fluorescence intensities along the diameter (y-axis) at the starting angle, 0°, over time (x-axis). The image in panel A represents t = 8 min. (*) represents the time point in C and D.
**E, F.** The intensity measurements at each point of the diameter are averaged over all angles, i.e. over the entire circumference at each distance from the center for each time point. Since the diameter sweeps across each point twice (once with each radius), this results in a symmetrical representation, of which only one half is shown in the final kymograph.
**F.** Resulting two-dimensional (kymograph) of calcium signaling in time and space. The vertical cyan lines in the graph represent the calcium signal in periderm and basal cells with the distance from the central cell represented along the y-axis.

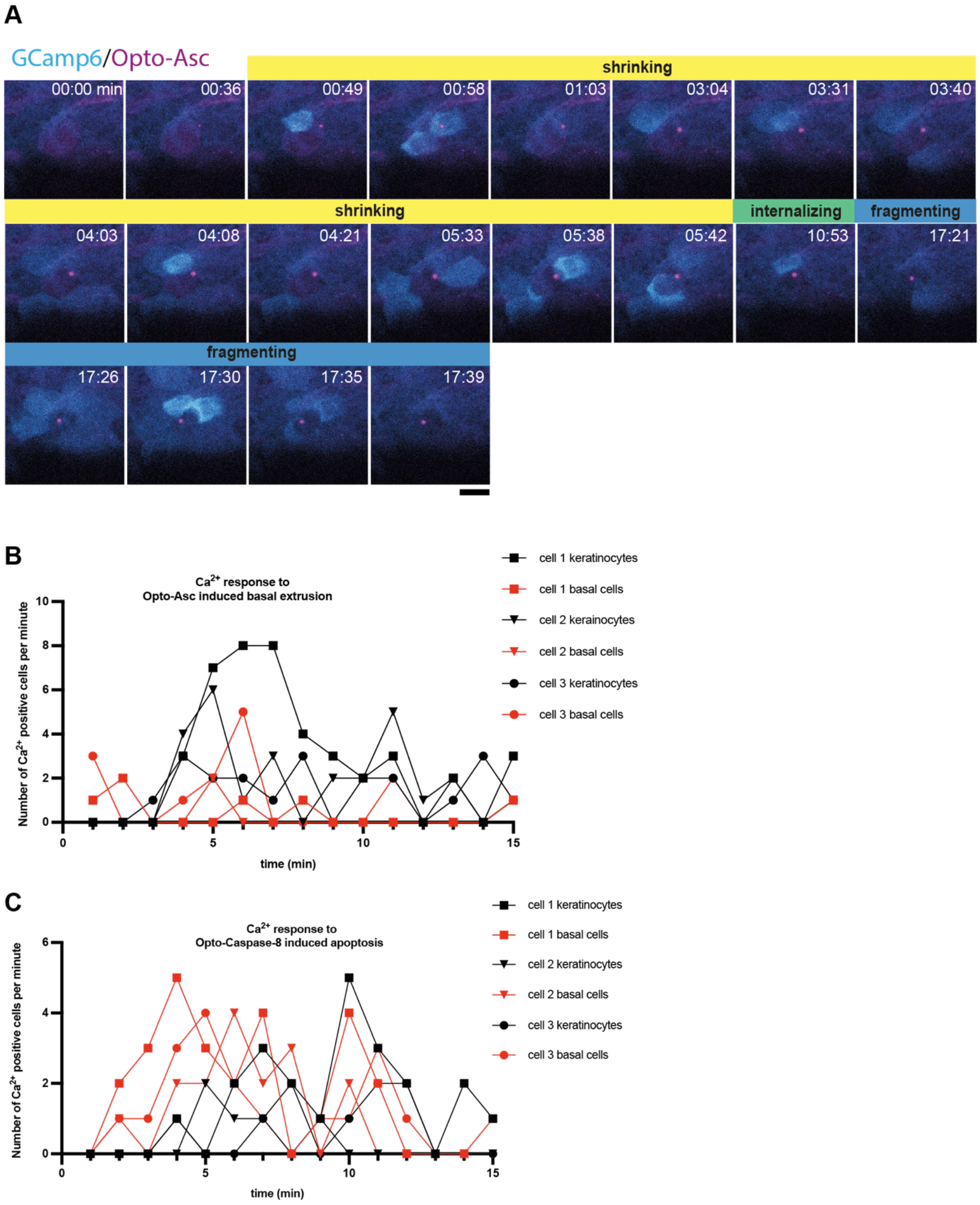


**Supplementary figure 7: $Ca^{2+}$ response of neighboring cells to Opto-Asc induced basal extrusion and its quantification.**

**A.** Example of $Ca^{2+}$ signaling in epithelial cells for basally extruding periderm cells imaged at high temporal resolution (15 frames per minute).

Opto-Asc (magenta) is induced in a periderm cell in the center of the frame by blue light (488nm). Both the speck and the $Ca^{2+}$ signal in the neighboring cell appear simultaneously at 00:49 min. The stages of cell death are indicated as defined in Figure 5: shrinking in yellow, internalizing in green, and fragmentation in blue.

**B-C.** $Ca^{2+}$ responses in cells adjacent to the dying cell.

Number of neighboring cells that show a $Ca^{2+}$ elevation in response to Opto-Asc-induced basal extrusion and Opto-Caspase-8-induced apoptosis. Three movies, indicated by triangles, squares and circles, were analyzed in each experiment.
Number of 1st neighbors (cells directly in contact with the dying cell) to Opto-Asc-induced dying cells (**B**) and to Opto-Caspase-8-induced dying cells (**C**) plotted over time. periderm cells are represented in black, and basal cells are in red.

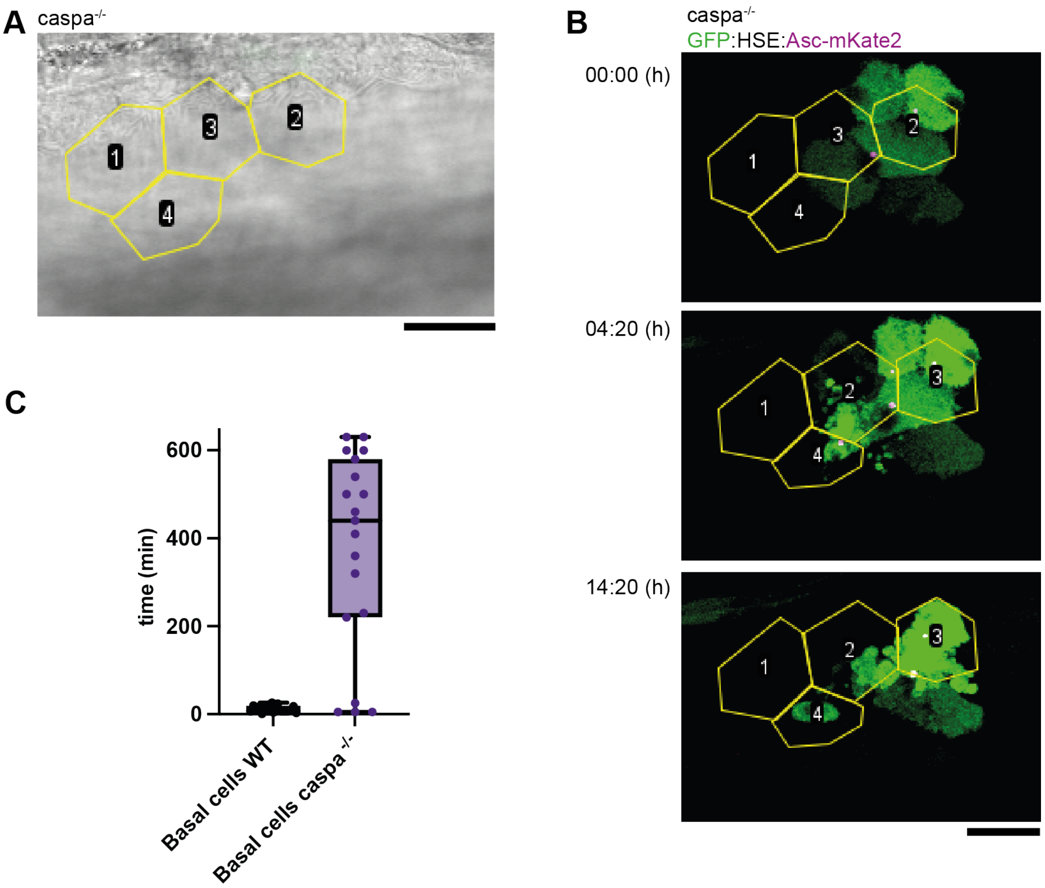


**Supplementary figure 8:**

**A.** Bright field image of the skin of a caspa-/- larvae showing the outlines of periderm cells (yellow) which were identified by scanning though the z-planes of the stack. Scale bar is 20 µm.

**B.** Time laps of basal cells transiently expressing *HSE: GFP/Asc-mKate2* underlying the periderm cells identified in **A** and outlines in yellow. GFP is in green, Asc.mKate2 in magenta. Scale bar is 20 µm.

**C.** Time between speck formation and cell death measure in minutes in basal cells of larvae of the *tg(HSE:Opto-Asc)* lines compared to basal cells in transiently expressing caspa$^{-/-}$ larvae.
